# Education can Reduce Health Disparities Related to Genetic Risk of Obesity: Evidence from a British Reform

**DOI:** 10.1101/260463

**Authors:** Silvia H. Barcellos, Leandro S. Carvalho, Patrick Turley

## Abstract

This paper investigates whether genetic makeup moderates the effects of education on health. Low statistical power and endogenous measures of environment have been obstacles to the credible estimation of such gene-by-environment interactions. We overcome these obstacles by combining a natural experiment that generated variation in secondary education with polygenic scores for a quarter million individuals. The additional schooling affected body size, lung function, and blood pressure in middle age. The improvements in body size and lung function were larger for individuals with high genetic predisposition to obesity. As a result, education reduced the gap in unhealthy body size between those with high and low genetic risk of obesity from 20 to 6 percentage points.

---

Educational policies may increase or decrease health disparities depending on whether they reinforce or counteract gene-related health disparities *(1)*. Both childhood circumstances, such as education, and genetic factors are independently associated with later life health (2,3,4). A growing literature suggests that health may also depend on the interaction between these two factors (5,6,7). Where strong gene-by-environment (GxE) interactions exist, modest effects of education on average may conceal larger effects for populations with particular genotypes and lead to underestimates of the benefits of schooling. We investigate this possibility by testing whether genetic makeup moderates the effect of an additional year of secondary education on middle age health.

After the publication of high-impact GxE studies *(8,9,10)*, controversies over the replicability of results tempered the enthusiasm for this research program. Low statistical power and endogenous measures of environment are believed to be main reasons for the limited replicability (11,12,13). Many GxE studies are low-powered because behavioral traits tend to be polygenic, meaning that they are influenced by a large number of genetic markers, each with a very small effect (14). Furthermore, the effect size of interactions is typically lower than that of direct effects (11). As a result, much of the previous literature, which focused on individual candidate genes (exceptions include *(15, 16, 17)*), was underpowered *(12).*

In addition, endogenous measures of environment may lead to biased estimates of GxE interactions *(18,19).* Measures of environment are “endogenous” when the outcome affects the environment (i.e., “reverse causality”) or when the relationship between the environment and the outcome is confounded by omitted third factors. Endogenous measures are a concern in our context because health in childhood may affect educational attainment, or self-control may drive both schooling decisions and health behaviors.

We overcome these obstacles by combining a natural experiment with polygenic scores (PGSs), which are indices constructed from millions of genetic markers. The natural experiment, a well-known compulsory schooling age reform in the UK, generated as-good-as-random variation in education: we estimate that 14 percent of students completed an additional year of secondary education as a result of this reform *(20,21).* The combination of this experiment with the use of PGSs—instead of a candidate-gene approach—for a sample of a quarter-million individuals make our analyses appropriately powered *(22)*.

Prior to the release of the complete genetic data used in this study, we wrote a comprehensive pre-analysis plan describing the construction of all variables to be used and the specification of all analyses to be run (see ref *22* and Appendix A). We strictly follow this plan below. Our plan was informed by our previous work, which used the same natural experiment and non-genetic data to estimate the effect of education on middle age health *(23).* We found that education reduced body size, improved lung function, and increased blood pressure, but that these effects were concentrated at specific parts of the health distribution (see Appendix C).

Following our previous work, we use UK Biobank (UKB) data (24) to study the same three health dimensions: body size, lung function, and blood pressure. To reduce concerns about multiple-hypothesis testing, we construct an index that is a weighted average of objective outcomes measuring each dimension *(25)*. The body size index includes body mass index (BMI), body fat percentage, and waist-hip ratio. The lung function index includes three spirometry measures. The blood pressure index includes multiple diastolic and systolic blood pressure measurements. We also construct a summary index that is a weighted average of the body size, lung function, and blood pressure indices (see Appendix A, section VI). We orient all four indices so a higher number corresponds to worse health. For each index, we study two types of outcomes: the continuous index measure and an indicator for whether the index is above a threshold specified in our pre-analysis plan *(22)*. These thresholds correspond to the values where we estimated the largest distributional effects in our previous work *(23)*. Although selecting the threshold this way leads to upward biased estimates of the effect of education, it does not lead to biased estimates of the GxE interaction (Appendix A, section XI). In Appendix I, we show our results are robust to alternative thresholds.

We construct PGSs for two traits for which large genome-wide association studies (GWAS) are publicly available: BMI *(26)* and Educational Attainment (EA) *(27)*. Individuals with higher genetic predisposition to EA might benefit more from an additional year of schooling because, among other reasons, the EA PGS is genetically correlated with cognitive performance *(27)*. For example, those with a higher EA PGS might receive higher incomes and work in better occupations as a result of the additional schooling, with potentially larger consequences for their health. Currently, there are no publicly available, sufficiently predictive GWAS for traits related to lung function and blood pressure. We opt therefore to investigate whether the BMI PGS moderates the effects of education on lung function and on blood pressure because BMI is genetically correlated with smoking and with coronary artery disease *(28)*. The PGSs were normalized to have mean zero and standard deviation one, and oriented so each PGS is positively correlated with its corresponding outcome *(29)*.The correlation between these two PGSs is −0.24.

Figure 1 shows health disparities between those with lower and higher genetic risk of obesity. It plots the fraction of study participants in the bottom, middle, and top terciles of the BMI PGS distribution with a health index above its corresponding threshold. To facilitate the comparison with Figure 3 estimates, we restrict the sample to participants who were born before September 1, 1957 and who dropped out before age 16. While 11% of those in the bottom PGS tercile had a body size above the threshold, this fraction was almost 3 times larger (31%) among those in the top tercile. The figure shows the BMI PGS is more predictive of the body size (*R*^2^ = 0.049) and summary indices (*R*^2^ = 0.021) than of the lung function (*R*^2^ = 0.002) and blood pressure indices (*R*^2^ = 0.002). See Appendix D for the corresponding figures for continuous outcomes and for predictive power of EA PGS.

**Fig. 1.**
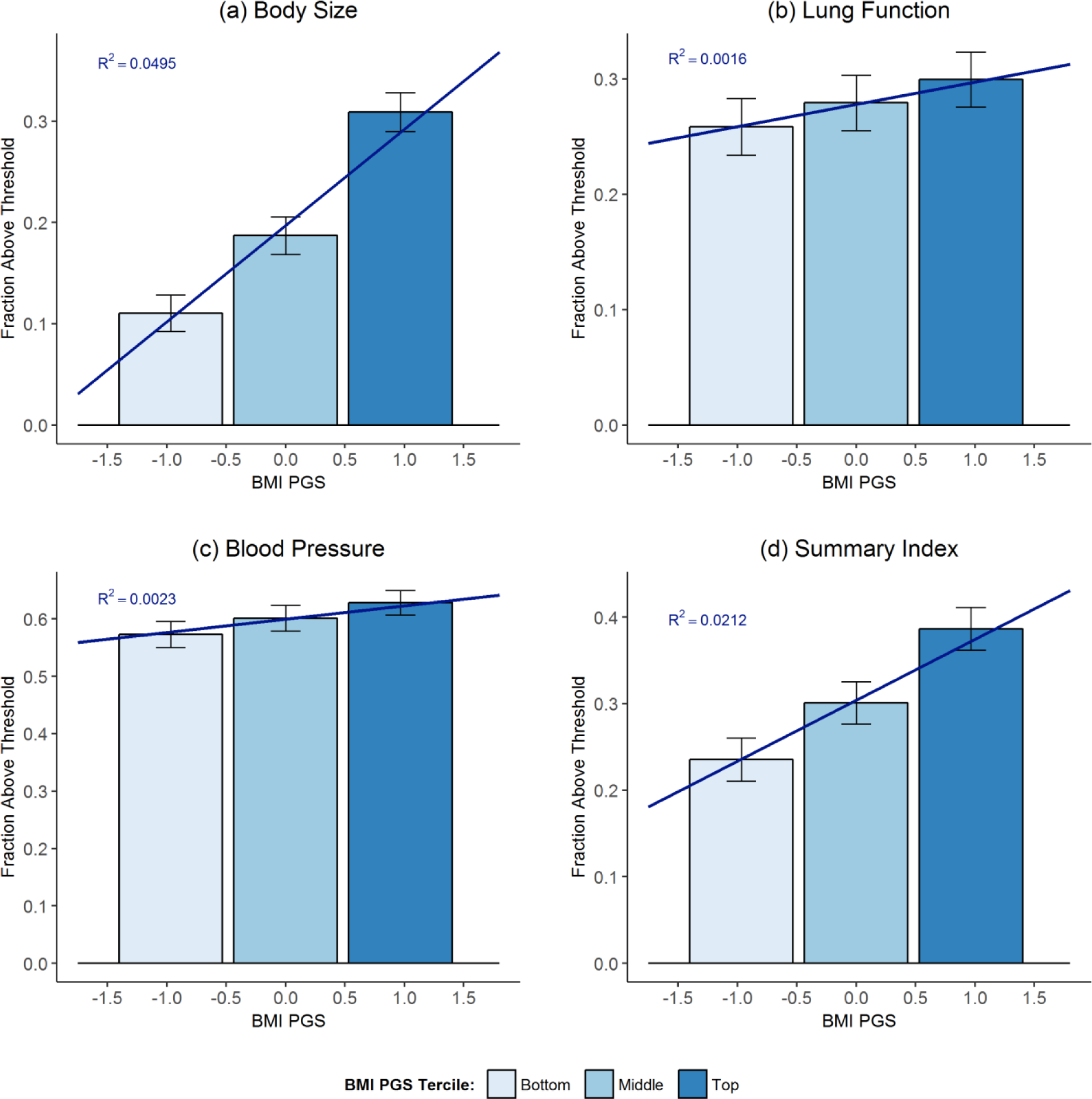
Health disparities by BMI PGS. Bars show means of binary measures of **(a)** body size, lung function, **(c)** blood pressure, and **(d)** summary indices for the bottom, middle, and top terciles of the BMI PGS distribution with 95% confidence intervals. Bars are centered at the median PGS value in the tercile. Sloped lines give linear projection of outcomes on BMI PGS. R^2^ gives the fraction of the variation in the outcome explained by the BMI PGS. To make estimates comparable to our estimates in Fig. 3, we restrict the sample to participants who were born before September 1, 1957 and who dropped out before age 16 and control for a quadratic polynomial in date of birth.

In 1972, England, Scotland, and Wales increased the minimum age at which students could drop out of school from 15 to 16 years. The reform affected only students born on or after September 1, 1957, generating a discontinuity in the relationship between education and date of birth.

Figure 2a shows that the fraction staying in school until age 16 increased *discontinuously* for those born after September 1, 1957. About 83% of those born between September 1956 and August 1957 stayed in school until at least age 16. This fraction is close to 97% among those born between September 1957 and August 1958, the first birth cohort affected by the reform. One can interpret this discontinuous change, which has been documented by previous studies *(21,30)*, as the effect of the reform on education. In the UKB sample, we estimate the policy increased the fraction staying in school until age 16 by 14 percentage points (see Appendix E). In our previous work, we showed that the policy also led individuals to obtain more qualifications, earn higher income, and work on occupations with higher socioeconomic status *(23)*.

**Fig. 2.**
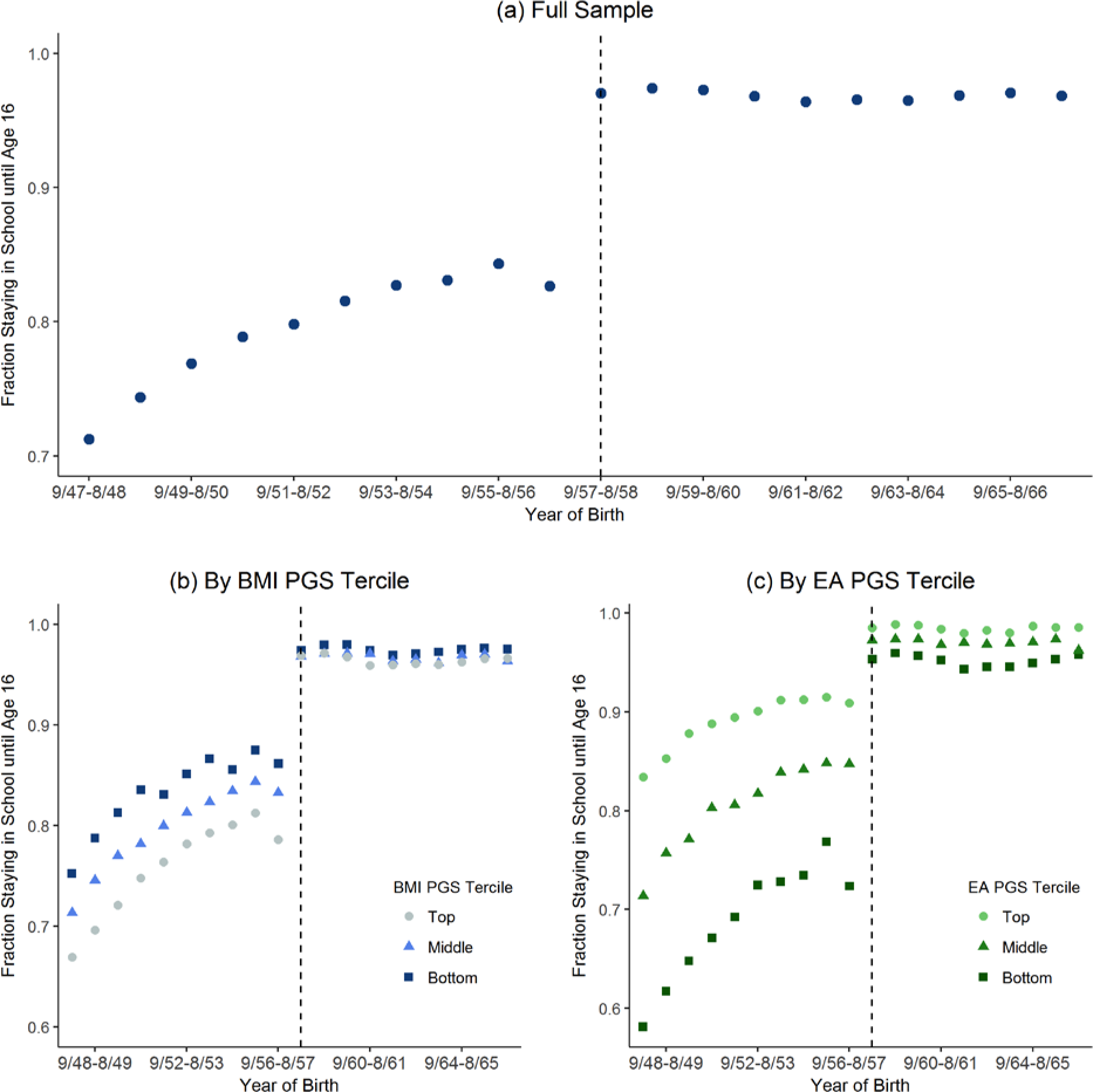
Fraction staying in school until age 16 by year of birth for **(a)** full sample, **(b)** bottom, middle, and top terciles of the BMI PGS distribution, and **(c)** bottom, middle, and top terciles of the EA PGS distribution. Dashed vertical lines mark the first birth cohort affected by the raising of the school-leaving age from 15 to 16.

To estimate the causal effect of education on health, we use a regression discontinuity design (RDD). Intuitively, the RDD compares the health outcomes of individuals born just before and just after September 1, 1957, controlling for cohort trends. Under the assumption that no other factors changed discontinuously at this birth date, any differences in health between these two groups born just days apart can be attributed to the causal effect of education. We offer evidence that this assumption holds in Appendix B. For example, we show that individuals born before and after this date are genetically similar. Genetic markers are useful to test the RDD assumption because genotypes are objectively measured, determined at conception, and immutable.

To investigate whether the effect of education on health varies with genetic makeup, we compare the discontinuous changes in health of groups with different PGSs, accounting for the differences in the fraction of individuals affected by the reform in different PGS groups. Figures 2a and 2b show that, among cohorts born before September 1957, those with *higher* BMI PGSs and those with *lower* EA PGSs were less likely to stay in school until age 16. Because all students were required to stay in school until age 16, the reform had a larger effect on those groups.

Formally, we estimate the following regression:

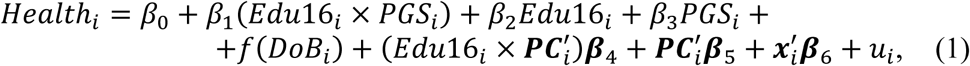

where *Health_i_* is a health index; *Edu16_i_* is an indicator for staying in school until age 16; *PGS_i_* is the BMI or EA PGS; *f(DoB_i_)* is a quadratic polynomial in date of birth (we allow for different pre- and post-trends); ***PC**_i_* is a vector of the first 15 principal components of the genotypic data; and ***x**_i_* is a vector of predetermined characteristics - namely age, age-squared, gender, month and country of birth. We include *Edu* 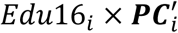 and 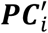 to correct for population stratification *(31,32)*. To account for the endogeneity of *Edu16_i_* and for the differential impacts of the reform on the education of groups with different PGSs, we estimate equation (1) through two-stages least squares (2SLS), using the reform as an instrument. We restrict the sample to participants born within 10 years of September 1, 1957 (*N* = 253,715). In Appendix H, we show our results are robust to tighter bandwidths and to linear trends.

Table 1 summarizes the main results (see Appendices E and G for additional results). We find that overall the effects of education on health depend on the BMI PGS. In 5 out of 8 regressions, the p-value on β_1_ is less than 0.05. In 2 cases, it is less than the Bonferroni-corrected value 0.05/16 = 0.0031. In contrast, there is no evidence that the effects of education on health depend on the EA PGS. None of the 8 regressions have p-values on the interaction term less than 0.05.

**Table 1.** Effect of staying in school until age 16 on health. 2SLS estimates. Above Threshold is an indicator of whether the health index is greater than the threshold specified in ***(Error! Bookmark not defined.).*** Edu16 is an indicator for staying in school until age 16 and is instrumented by an indicator for being born after September 1, 1957. The “P-value for H_0_: No Effect of Education” is the p-value from a joint test that *β_1_ = β_2_* = 0. The last row shows means of the dependent variable among pre-reform compliers, defined as individuals born before September 1, 1957 who dropped out before age 16.

|  | Continuous |  |  |  | Above Threshold |  |  |  |
| --- | --- | --- | --- | --- | --- | --- | --- | --- |
|  | Body Size | Lung Function | Blood Pressure | Summary Index | Body Size | Lung Function | Blood Pressure | Summary Index |
| **Interaction with BMI PGS** |  |  |  |  |  |  |  |  |
| BMI PGS × Edu16 | 0.028<br>(0.033) | -0.091**<br>(0.044) | 0.073**<br>(0.036) | -0.010<br>(0.042) | -0.057***<br>(0.011) | -0.037**<br>(0.016) | 0.016<br>(0.018) | -0.101***<br>(0.016) |
| Edu16 | -0.119<br>(0.096) | -0.147<br>(0.124) | 0.118<br>(0.102) | -0.092<br>(0.122) | -0.060*<br>(0.035) | -0.089*<br>(0.048) | 0.106**<br>(0.051) | -0.058<br>(0.050) |
| BMI PGS | 0.263***<br>(0.030) | 0.127***<br>(0.040) | 0.024<br>(0.033) | 0.206***<br>(0.038) | 0.124***<br>(0.010) | 0.048***<br>(0.015) | 0.025<br>(0.016) | 0.152***<br>(0.015) |
| P-value for $H_0$: No Effect of Education | 0.455 | 0.002 | 0.002 | 0.545 | $7.97 \times 10^{-10}$ | $6.09 \times 10^{-5}$ | 0.004 | $1.50 \times 10^{-14}$ |
| **Interaction with EA PGS** |  |  |  |  |  |  |  |  |
| EA PGS × Edu16 | -0.093<br>(0.059) | -0.001<br>(0.074) | -0.024<br>(0.060) | -0.054<br>(0.074) | -0.013<br>(0.020) | 0.030<br>(0.028) | 0.021<br>(0.030) | 0.049*<br>(0.029) |
| Edu16 | -0.192<br>(0.119) | -0.172<br>(0.148) | 0.117<br>(0.121) | -0.146<br>(0.147) | -0.089**<br>(0.042) | -0.085<br>(0.056) | 0.117*<br>(0.060) | -0.067<br>(0.059) |
| EA PGS | -0.036<br>(0.057) | -0.060<br>(0.072) | -0.054<br>(0.058) | -0.076<br>(0.072) | -0.013<br>(0.020) | -0.043<br>(0.027) | -0.054*<br>(0.029) | -0.078***<br>(0.028) |
| P-value for $H_0$: No Effect of Education | 0.257 | 0.037 | 0.016 | 0.576 | 0.006 | $3.36 \times 10^{-7}$ | 0.014 | $2.11 \times 10^{-8}$ |
| Observations | 249,699 | 203,048 | 253,377 | 200,398 | 249,699 | 203,048 | 253,377 | 200,398 |
| Dependent Variable Mean among Compliers | 0.261 | 0.269 | 0.090 | 0.311 | 0.215 | 0.287 | 0.611 | 0.317 |

We can reject the hypothesis that staying in school until age 16 has no effects on health in middle age. In 12 out of 16 regressions, the p-value of the joint test that *β*_1_= *β_2_* = 0 is less than 0. 05 and in 10 cases less than the Bonferroni-corrected value of 0.05/16 = 0.0031. The direction of these results is consistent with previous work *(23,30)*.

For the binary measures of the body size, lung function, and summary indices, the improvements in health are larger for individuals with higher BMI PGSs. Similarly, for the continuous measure of lung function, improvements in health are larger for individuals with higher BMI PGSs. While the estimate for the continuous measure of blood pressure suggests an interaction of the BMI PGS and education, there are reasons to question the credibility of this particular result: its marginal significance (p-value of 0.041), the weak direct effect of the PGS (p-value of 0.458), and the low power anticipated in the pre-analysis plan (in the most optimistic case, 17% power to detect an effect at 5% significance).

The results shown in Table 1 assume that the effect of staying in school until age 16 varies linearly with the PGS. In Figure 3 we adopt a more nonparametric specification and estimate separate effects for the bottom, middle, and top terciles of the BMI PGS distribution. The bars show point estimates of the effects on the binary outcomes with 95% confidence intervals. Figures presented in Appendix F for the continuous measures and EA PGS show results qualitatively similar to the corresponding results in Table 1.

**Fig. 3.**
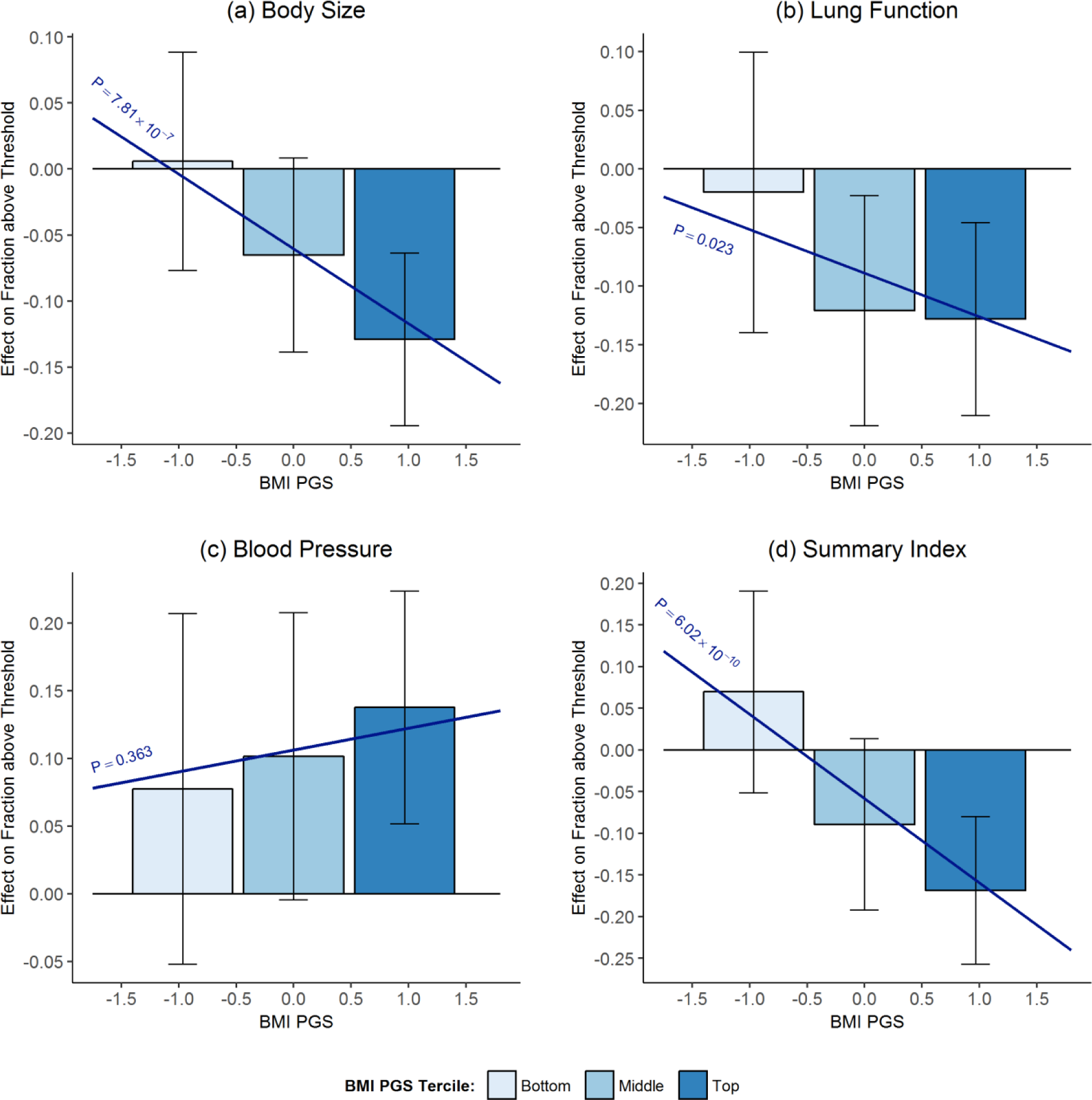
Does the effect of staying in school until age 16 depend on the BMI PGS? Bars show 2SLS point estimates of effect of staying in school until age 16 on binary measures of **(a)** body size **(b)** lung function **(c)** blood pressure and **(d)** summary indices for the bottom, middle, and top terciles of the BMI PGS distribution. Bars are centered at the median PGS value in the tercile. Brackets show 95% confidence intervals. Sloped lines plot 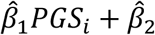. “P” corresponds to the p-value of *H_0_: β_2_* = 0.

Figure 3 shows that education reduced the health disparities in body size shown in Figure 1. For the top tercile of the BMI PGS distribution, staying in school until age 16 reduced the fraction above the body size threshold by 13 percentage points. For the bottom tercile, there was a modest, statistically insignificant increase. As a result, the additional year of education reduced the gap in “unhealthy body size” (i.e., being above the body size threshold) between the top and the bottom PGS terciles from 20 to 6 percentage points. To put this into perspective, individuals at the threshold have an average BMI of 31.4—the medical cutoff for obesity is 30. In Appendix J, we show results for medical cutoffs for BMI and blood pressure.

Our results challenge the notion of genetic determinism; yet the question of why we observe larger health improvements for those with higher genetic predisposition to obesity remains. One hypothesis is that the additional year of schooling led those with higher genetic risk to make larger adjustments to their health behaviors. We find no evidence to support this hypothesis when using UKB data on diet and physical activity (see Appendix K), but we stress that these data have several important limitations (e.g., data available only for part of the sample; self-reported diet measures). An alternative hypothesis is that a change in health behaviors of a given magnitude translates into larger health improvements for those with higher genetic risk. We leave to future work to investigate these hypotheses.

Investigating the generalizability of our results will be an important next step. As the availability of genotypic data linked to social surveys grows and allows for larger GWAS, new PGSs with reasonable predictive power will become available for a variety of health and behavioral phenotypes. Reproducing the analyses above with different PGSs and in different contexts will increase our understanding of how environmental and genetic factors interact and of the role social policy can have in mitigating possible inequalities arising from genetic background.

## Online Appendix

## APPENDIX A Pre-Analysis Plan

This appendix shows the pre-analysis plan registered before the release of the full UKB genetic data, which can also be found at osf.io/9dyfz

In the version below, we have corrected some ambiguous notation and harmonized the terminology to match that of the body of the paper. We have added comments boxes to highlight each instance where the pre-registered plan and the plan below deviate. We also use text boxes to highlight which contingency was taken in the case that possible alternative decisions could be made.

### I. Background

Studies about the causal effect of education on health have focused on average effects (e.g., Lleras-Muney 2005; Clark and Royer 2013), although there is some evidence that educational interventions may have larger impacts on the later-life health of some individuals than of others (Barcellos et al. 2017). Here we propose a follow-up to Barcellos et al. (2017), in which we examine whether genetic factors play a role in this heterogeneity, and more precisely, if genetic predictors of health and education are able to identify those for whom education has the largest effect on health later in life.

### II. The 1972 ROSLA

This research will use a regression discontinuity design to study the heterogeneous effects of education on health. In 1972 England, Scotland, and Wales raised their minimum school-leaving age from 15 to 16 for students born on or after September 1, 1957 (students born before this date could drop out at age 15), generating a discontinuity in the relationship between education and date of birth at the September 1, 1957 “cutoff.” There are a number of studies that have exploited changes in compulsory schooling laws to study the causal effect of education on average health (e.g., Lleras-Muney 2005; Albouy and Lequien 2009; Silles 2009; Powdthavee 2010; Kemptner et al. 2011; Jürges et al. 2013) and several that have considered the 1972 ROSLA reform in particular (Clark & Royer 2013, Davies et al. 2017, Barcellos et al. 2017). This research builds on that literature by measuring how the causal effect of education on health varies by a person’s genetic risk for poor health and genetic risk for educational achievement.

### III. Data

We will use data from the UK Biobank, a large, population-based prospective study initiated by the UK National Health Service (NHS) (Sudlow et al. 2015). Between 2006 and 2010, invitations were mailed to 9.2 million people between the ages of 40 and 69 who were registered with the NHS and lived up to about 25 miles from one of 22 study assessment centers distributed throughout the UK (Allen et al. 2012).^1^ The sample is composed of 503,325 individuals who agreed to participate (i.e., a response rate of 5.47%). Although the sample is not nationally representative, our estimates have internal validity because there is no differential selection on the two sides of the September 1, 1957 cutoff (see Barcellos et al. 2017 and Davies et al. 2017).

Study participants went through an assessment that comprised a self-completed touch-screen questionnaire; a brief computer-assisted interview; physical and functional measures; and collection of blood, urine, and saliva. The collection of physical measures which we use in our analysis included anthropometrics, spirometry, and blood pressure and was standardized across centers. These measures were gathered by trained nurses or healthcare practitioners. About 100,000 participants also wore accelerometers that recorded physical activity for 7 days. Every participant was genotyped.

Our health measures will be based on the four objective health indexes studied by Barcellos et al. (2017): a body size index, a lung function index, a blood pressure index, and a summary index composed of the other three indexes. If other objective measures that are not currently available become available over the course of this project (e.g., blood glucose levels), we may study those in addition to these indexes.

In the pre-analysis plan we used the terms “anthropometrics index” and “body size index” interchangeably. We also used the terms “spirometry index” and “lung function index” interchangeably. To make it consistent with the body of the paper, we opted for replacing “anthropometrics index” with “body size index” and for replacing “spirometry index” with “lung function index.” For the same reason, we opted for replacing “general health index” with “summary index”, which is the terminology used in the body of the paper.

A description of how the indexes used in Barcellos et al. (2017) were constructed is found below.

### Body Size

An anthropometric index will be constructed from three measures: body mass index (BMI), body fat percentage, and waist-hip ratio.^2^ A bioimpedance analyzer was used to calculate body fat percentage. This device passes a low electrical current through the body. Water conducts electricity. While fat contains very little water, muscle contains 70% water. The bioimpedance analyzer calculates body fat from the speed of the current: The slower the signal travels, the greater the fat content.

### Lung Function

A spirometry test was conducted to measure participants’ lung function. The spirometer is a small machine attached to a mouthpiece by a cable that measures the volume and speed of air after a forced exhale. Participants were asked to fill their lungs as much as possible and to blow air out as hard and as fast as possible in the mouthpiece.^3^ Three parameters were measured: 1) forced expiratory volume in the first second is the amount of air exhaled during the first second; 2) forced vital capacity is the total amount of air exhaled during the forced breath; and 3) peak expiratory flow is the fastest rate of exhalation. These parameters are used to assess pulmonary conditions, such as chronic obstructive pulmonary disease and asthma. We follow DeMateis et al. (2016)’s criteria to identify acceptable expiratory maneuvers in the UK Biobank data. Valid spirometry measures are available for 79% of our sample.

### Blood Pressure

Two measurements were taken of the diastolic and systolic blood pressures of each study participant. We will use the average of these two measurements.

### Summary Indexes

In order to reduce the number of outcomes and partly address concerns about multiple hypothesis testing, we will construct for each of three health dimensions a summary index that is a weighted average of the different outcomes measuring that dimension:

<u>Body size</u>: body mass index, waist-to-hip ratio, and body fat percentage;

<u>Lung function</u>: forced expiratory volume in the first second, forced vital capacity, and peak expiratory flow;

<u>Blood pressure</u>: diastolic and systolic blood pressures.

Before combining these sub-measures into the three indexes, each measure will first be standardized separately by gender, using as a reference those born in the 12 months before September 1, 1957. We then will combine these measures into the three indexes following the procedure proposed by Anderson (2008), an approach that assigns weights based on the variance-covariance matrix of the input measures. Finally, we will construct a fourth <u>summary index</u> that is a summary of the body size, the lung function, and the blood pressure indices, using the same weighting procedure. We will construct all four indices so that a higher number corresponds to worse health.

In order to preserve the cutoffs derived in our previous work (see section VI. Health Outcomes), we standardized the measures using the entire sample used in Barcellos et al. (2017), including those without valid genotypic data and non-whites. After the standardization, we drop nonwhites and those without genotypic data or whose genotypic data did not pass quality controls.

### IV. Polygenic Scores

Genetic heterogeneity will be measured using polygenic scores that predict poor health and educational attainment. These scores are a weighted sum of single-nucleotide polymorphism (SNP) genotypes:

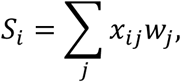

where *S_i_* is the genetic predictor of a trait (such as BMI or education) for individual *i, x_i,j_* ϵ {0,1,2} is a count of the number of reference alleles for individual *i* at SNP *j*, and *w_j_* is a weight associated with SNP *j.*

In the pre-analysis plan we made a notation mistake and wrote that “*S_i,j_* is the genetic predictor of trait *j*”, which was confusing because *j* was also used to index the SNP. To correct this mistake we dropped the subscript *j* from *S_i,j_*.

The weights used in a polygenic score are based on genome-wide association study (GWAS) summary statistics. A GWAS is a series of regressions of some outcome onto the genotype of a single SNP and a set of covariates. These covariates normally include sex, age, and the first several principal components of the genetic data. The principal components are included to control for omitted variable bias that may occur as a result of an individual’s ancestry (Price et al. 2006).

GWAS summary statistics for a number of traits are publicly available. To produce the final weights, we transform these summary statistics using a standard method, LDpred (Vilhjalmsson 2015). In addition to GWAS summary statistics, LDpred requires a reference sample to estimate the linkage disequilibrium structure (i.e., the correlation structure) of the genotypic data and an assumption about the fraction of SNPs which are truly associated with the outcome. Following the procedure described in Turley et al. (2017), we will use data from the 1000 Genomes Project as a reference sample and will assume that 100% of SNPs are associated with the outcome of interest. This assumption is unlikely true, but varying this parameter usually has little effect on the predictive power of polygenic scores produced by LDpred.

The two polygenic scores we plan to use for this study are for body-mass index (BMI) and educational attainment (EA). These were chosen due to their relevance to the outcomes considered and because large GWASs for these traits are publicly available: Locke et al. (2015) for BMI and Okbay et al. (2016) for EA. In the Health and Retirement Study, a score based on the GWAS results from Locke et al. (2015) explains 7% of the variation in BMI and a score based on Okbay et al (2016) explains roughly 5% of the variation in educational attainment. If larger-scale GWASs for these traits become available while this research is still in progress, we will use those instead. Additionally, if polygenic scores become available for traits related to our indexes (e.g., smoking behavior or blood pressure) that explain more than 5% of the variation in our health indexes in the UKB, we will also use those.

To avoid overfitting, it is important that the sample used for estimating the weights does not include individuals from the prediction sample. While neither Locke et al. nor Okbay et al. use data from the UKB, if new GWAS summary statistics become available that include the UKB, we will request results that omit it.

We note, however, that we can use the UKB data to augment the published GWASs in two ways that won’t lead to bias due to over-fitting. First, for each outcome that is measured in the UKB and for which we have a polygenic score, we will run a GWAS of that outcome using individuals in the UKB that are not included in any of the main analyses. (For example, this includes those born more than 10 years from the September 1, 1957 threshold and those not born in England, Scotland, and Wales.) We will follow the quality control protocols described in the Supplementary Note of Turley et al. (2017) to conduct this GWAS. The UKB GWAS results will then be meta-analyzed with the existing published GWAS results for the outcome.

Second, we will use a cross-validation-style procedure to use even more of the sample to boost power in the polygenic score. Specifically, we will hold out 10% of the sample and include the remaining 90% in the GWAS in the UKB described above. After meta-analyzing the UKB GWAS with the published GWAS results, we will create a polygenic score for the held-out 10% using LDpred. We will repeat this 10 times for each possible hold-out sample

This approach will induce some correlation in the polygenic scores between individuals, however, that would affect the standard errors. There is not a well-developed literature on how exactly the standard errors should be corrected. A variation on how this may be done is found in Chernozhukov et al. (2017), though it would need to be adapted to apply to the cross-validation procedure described here. If we cannot adapt this method to obtain valid standard errors or identify one that is more appropriate, we will forego this cross-validation procedure, restricting the GWAS sample to that used in the published work and the individuals in the UKB that are not included in the regression discontinuity analysis as described above.

We forewent the cross-validation-style procedure because we could not adapt this approach to obtain valid standard errors.

### V. The Regression Discontinuity (RD) Model

We would like to estimate the causal model

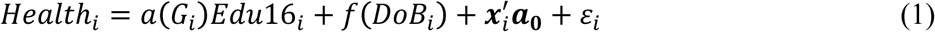

where *Health_i_* is some health outcome of interest for individual *i* (such as the body size index defined above), *G_i_* is that person’s genotype, *Edu16_i_* is an indicator of whether the individual stayed in school until at least age 16, *f* (*DoB_i_*) is a term that captures trends in health before and after the ROSLA reform, 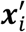 is a set of controls (including a constant and the genetic variables), and ε_*i*_ is the residual. We will use the same controls as in Barcellos et al. (2017). Notice that the causal effect of education on the health outcome is written as a function of *G_i_*, allowing this effect to capture genetic heterogeneity. We will describe our choice of *a* (*G_i_*) in the following section.

Of course, a person’s educational decisions may be correlated with factors in the residual, which would result in bias if this model were estimated directly. Following the literature, we will use the 1972 ROSLA as an instrument for *Edu16_i_* and estimate this model using a regression discontinuity (RD) design (see Barcellos et al. 2017).

For our first stage regression (i.e., the effect of the reform on *Edu16_t_*), we estimate the model

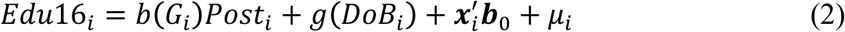

where *Post_i_* is an indicator variable for whether the individual was born on or after 1 September 1957, which is the threshold determining whether a person had to stay in school until age 16 according to the 1972 ROSLA. The corresponding reduced-form regression would be

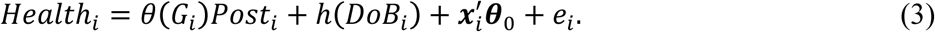

### VI. Health Outcomes

We will consider two types of health outcomes for this research. First, we will use the four indexes described in Section III. The results from analyses using these continuous outcomes can be interpreted as the mean effect of staying in school until at least age 16 on the outcome.

Second, we will construct binary outcomes from the indexes that equal one when the index is greater than some threshold. The binary outcomes may enable better powered analyses since Barcellos et al. (2017) find that the 1972 ROSLA had an impact on only part of the distribution for each of these indexes. We will therefore use the threshold at which the largest direct effect was estimated in Barcellos et al. (2017): 1.036 for the Body Size Index, 0.786 for the Lung Function Index, −0.215 for the Blood Pressure Index, and 0.786 for the (combined) Summary Index. We highlight that because we will use the same data for this research as was used in Barcellos et al. (2017), selecting the threshold in this way will lead to direct-effect estimates that are biased away from zero in expectation. We, however, are interested in the interaction term. It can be shown that the direct effect is uncorrelated with the interaction effect, so the bias in the direct effect should not affect the interaction coefficient estimates (see section XI. Bias of Interaction Coefficient). Nevertheless, as a robustness analysis for our main specifications, we will also perform our analysis using a number of thresholds across the distribution of the index.

### VII. Genetic Heterogeneity

As our main specification, we will define *a* (*G_i_*), *b* (*G_i_*), and θ(*G_i_*) as linear combinations of the scores being studied and the first 15 principal components of the genetic data.^4^ That is,

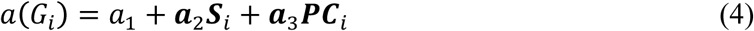

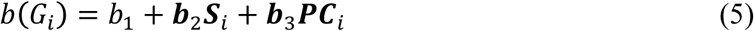

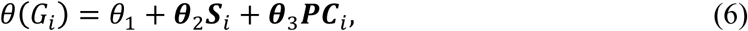

where ***S**_i_* is a vector of polygenic scores for individual *i* and ***PC**_i_* is a vector of the principal components of the genetic data for individual *i.* Using this specification, a_1_ + **a**_2_***S**_i_* may be thought of as the causal effect of an additional year of education for an individual with polygenic-score vector ***S**_i_*. As our main specifications, ***S**_i_* will be a scalar equal to either the BMI or EA polygenic score, but we will also perform secondary analyses where both polygenic scores are included in ***S**_i_* in order to test whether each polygenic score can explain any variation in the causal effect of education on health holding the other polygenic score constant.

As an additional, secondary, more nonparametric specification, we will create a set of indicator variables for each score of whether an individual has a polygenic score in the lowest, middle, or highest third of the sample. Then *a* (*G_i_*), *b*(G_*i*_), and θ(*G_i_*) will have the same functional form as before, but ***S**_i_* is a vector composed of the full set of binary variables defined above, omitting the lowest group. We chose to divide the data into three categories because at least 3 points are needed to observe a trend, but we would like to maximize the power of each coefficient by keeping the size of each group as large as possible.

### VIII. Specifications

To estimate this model by RD, we make several specification decisions. As in Barcellos et al. (2017), for our main specifications, we adopt a global polynomial approach in our estimation (see Lee and Lemieux 2010). We restrict the data to study participants born within 10 years of September 1957 - that is, born between September 1, 1947 and August 31, 1967 - and use a quadratic polynomial in date of birth to capture cohort trends.^5^ If possible, we will use date of birth measured in days, though we may be restricted to using month of birth due to data constraints. If we use month of birth, we will cluster our standard errors by month of birth. We use triangular kernel weights in all of our regressions.

We use date of birth measured in days and henceforth do not need to cluster the standard errors by month of birth. We calculate robust standard errors.

As controls, we use sex, age in days (at the time of the baseline assessment) and age squared, dummies for ethnicity, dummies for country of birth, and dummies for calendar month of birth (to control for seasonality).^6^

We do not use dummies for ethnicity as controls because everyone in our sample is white.

The first-stage regression (2) and the reduced form regression (3) will be estimated using ordinary least squares. Equation (1) will be estimated via two-stage least squares, where we will use the indicator variable for whether the individual was born on or after 1 September 1957, *Post_i_*, to instrument for whether the individual stayed in school until at least age 16, i.e., *Edu16_i_* (and the interactions of *Edu16_i_* with *G_i_* and ***PC**_i_* will be instrumented using the interactions of *Post_i_* with *G_i_* and ***PC**_i_*).

Of course, we will verify that all of our results are robust to deviations from each of these decisions. For example, we will test combinations of a number of shorter bandwidths, linear trends, and exclusion of control variables. Note that Barcellos et al. (2017) perform a McCrary test (McCrary 2008) and a number of balance tests to strengthen the evidence that the assumptions of a regression discontinuity design are met. We will be using the same data (albeit with an updated polygenic score), but we will replicate those results nonetheless.

### IX. Analysis of Channels

This project will also include several secondary analyses to better understand the potential channels through which any observed effects are found. These will consist primarily of analyses identical to those described above, replacing the health indexes with other outcomes in the UKB (as in Barcellos et al. 2017), including measures of household income, labor market outcomes, neighborhood deprivation, occupational prestige, diet, and smoking behavior. Since these analyses are only secondary we have not listed exhaustively all of the channels we will consider.

In Appendix K we investigate the channels through which the effects on health are moderated by genetic risk of obesity by examining whether those with higher genetic risk of obesity make larger adjustments to their diet and physical activity.

### X. Power Calculations

To calibrate our anticipated effect sizes for this power calculation, we look both to the literature and to preliminary results. There are many existing studies that consider interactions between environmental or policy variables and single SNP variables (see Duncan and Keller 2011 for a review), and a few that interact an endogenous environmental variable with a polygenic score (e.g., Belsky et al. 2016, Barth et al. 2017). There are few published studies, however, that estimate the effect of an interaction of a policy variable with a polygenic score (an exception is Schmitz and Conley 2016). This distinction is important since policy variables tend to explain much less of the variation in an outcome relative to endogenous environmental variables, and therefore larger samples are needed to be well powered to detect the direct and interaction effects. An advantage of using exogenous policy variables, however, is that the results of the analysis will have a causal interpretation.

In the studies listed above, the polygenic scores used in those studies explained around 5% of the variation in the outcome they study. For studies using endogenous environmental variables, the polygenic scores similarly explained roughly 5% of the variation. In contrast, the ROSLA, an exogenous policy variable, only explains around 0.01% of the variation in the health outcomes considered in Barcellos et al (2017). This suggests that the explanatory power of the interaction term in our setting (and therefore our power to detect a significant effect for a given sample size) may be substantially smaller than what has been observed in previous work.

Table 1 below reports the two-stage least squares estimates following a similar analysis plan to the one described above. These results were prepared as part of a lecture on the power of gene-environment interaction studies and therefore do not correspond exactly to what will be done in this research. The results, however, should give ballpark estimates of the power to detect significant interactions in the UKB. The key differences between how these results were obtained and analysis plan described above are: (i) these results are based on the preliminary release of the UKB data (roughly a quarter of the full sample), (ii) the health outcome measures only include the continuous indexes (omitting the binary measures), (iii) the polygenic scores were included in the model jointly rather than separately, and (iv) date of birth was measured in months, and hence the standard errors were clustered by month of birth. Note that few of the interaction results reported are statistically significant even at the 5% level.

**Table 1:** Preliminary Estimates

|  | Body<br>Size | Lung<br>Function | Blood<br>Pressure | Summary<br>Index |
| --- | --- | --- | --- | --- |
| Edu16 | -0.325<br>(0.298) | -0.055<br>(0.345) | -0.436<br>(0.315) | -0.291<br>(0.318) |
| BMI PGS | 0.133***<br>(0.047) | 0.145**<br>(0.062) | 0.036<br>(0.046) | 0.151***<br>(0.054) |
| EA PGS | 0.004<br>(0.059) | 0.003<br>(0.070) | 0.004<br>(0.063) | 0.026<br>(0.075) |
| BMI PGS * Edu16 | 0.089<sup>†</sup><br>(0.055) | -0.145**<br>(0.07) | 0.018<br>(0.052) | -0.022<br>(0.062) |
| EA PGS * Edu16 | -0.077<br>(0.066) | -0.051<br>(0.076) | -0.062<br>(0.069) | -0.112<br>(0.083) |

<sup>†</sup>, 0.15; \*, 0.1; \*\*, 0.05; \*\*\*, 0.01

To be consistent with the body of the paper, in Table 1 we use “PGS” to refer to the polygenic scores instead of “Score”, which was the term used in Table 1 of the pre-analysis plan.

We believe that many of these coefficients will be significant once the full data are available, however, for four reasons. First, the sample will be approximately four times larger, cutting the size of the standard errors in half. Second, with a more powerful polygenic score, the magnitude of the coefficients should rise. Third, the parallel specifications with binary variables may have larger effects since the binary variables correspond to regions of the outcome distribution more strongly affected by the 1972 ROSLA. Fourth, if date of birth is measured in days, our estimates should become more precise both because we can capture more of the variation with a more precise running variable and because it removes the need to cluster standard errors by month of birth. On the other hand, due to the winners’ curse, it may be that the coefficients estimated here are larger in magnitude that the true coefficients.

In Table 2, we report the power of detecting effects with a p-value of less than 0.05 assuming that the true effect is (a) half what is observed in our data, (b) exactly what is observed in our data, and (c) 20% larger than what is observed in our data. In all cases, we assume that the standard errors are half as large as what is reported in Table 1, assuming a four-fold increase in sample size. We model the aggregate effect of the other factors affecting power (e.g., using a more powerful polygenic score) by adjusting the assumed effect size, as described in this paragraph.

In the pessimistic case (a), we do not exceed 80% power for any of the interaction terms, though we have greater than 50% power for 2 of the 8 estimates. If the true effect is equal to the estimate in our preliminary analyses, we have 80% power to reject the null for these same two coefficients. In the optimistic case (c), 3 of the 8 estimates have 80% power, and one more has a power of 77%.

We highlight that, consistent with the results of Barcellos et al. (2017), the power remains low for the main effect (Edu16) even in the optimistic case, likely due to treatment effect heterogeneity.

**Table 2:** Power at *p* = 0.05

|  | Case | Body<br>Size | Lung<br>Function | Blood<br>Pressure | Summary<br>Index |
| --- | --- | --- | --- | --- | --- |
| Edu16 | a | 0.28 | 0.06 | 0.41 | 0.21 |
|  | b | 0.59 | 0.06 | 0.79 | 0.45 |
|  | c | 0.71 | 0.07 | 0.89 | 0.56 |
| BMI PGS | a | 0.94 | 0.83 | 0.17 | 0.94 |
|  | b | 1.00 | 1.00 | 0.35 | 1.00 |
|  | c | 1.00 | 1.00 | 0.44 | 1.00 |
| EA PGS | a | 0.05 | 0.05 | 0.05 | 0.07 |
|  | b | 0.05 | 0.05 | 0.05 | 0.11 |
|  | c | 0.05 | 0.05 | 0.05 | 0.13 |
| BMI PGS * Edu16 | a | 0.53 | 0.74 | 0.07 | 0.07 |
|  | b | 0.90 | 0.99 | 0.11 | 0.11 |
|  | c | 0.96 | 1.00 | 0.13 | 0.13 |
| EA PGS * Edu16 | a | 0.31 | 0.13 | 0.20 | 0.39 |
|  | b | 0.65 | 0.27 | 0.44 | 0.77 |
|  | c | 0.77 | 0.34 | 0.54 | 0.87 |

To be consistent with the body of the paper, in Table 2 we use “PGS” to refer to the polygenic scores instead of “Score”, which was the term used in Table 2 of the pre-analysis plan.

### XI. Bias of Interaction Coefficient

To avoid confusion, this section, which was titled “Appendix” in the pre-analysis plan, was renamed “Bias of Interaction Coefficient.”

To be consistent with the body of the paper, we use “Post” to refer to the indicator variable of being born on or after September 1, 1957 instead of “After*_i_*”, which was used in the preanalysis plan.

We will show that the estimation error in the coefficient associated with the *Post* term of the regression is uncorrelated with the estimation error in the coefficient associated with the interaction variable, *Post_i_ × S_i_*, where *S_i_* is the polygenic score of interest.

First note that the variance-covariance matrix of the estimation is proportional to **Ω**^−1^, where **Ω** is the variance-covariance matrix of the variables in the model, including the variable *Post_i_*, *S_i_*, their interaction, and the controls. By the assumption of the RD design, *Post_i_* is independent to *S_i_* and the controls. Furthermore,

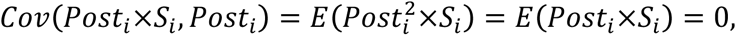

so *Post_i_* is uncorrelated with the interaction as well. This means that the elements of the row and column of **Ω** that corresponds to the variable *Post_i_* are uniformly zero. This also means that the elements of the row and column of **Ω**^−1^ that corresponds to the variable *Post_i_* are also uniformly zero. This implies that the estimation error of the coefficient for the *Post_i_* variable is uncorrelated with the estimation error of the coefficient of the interaction term.

## APPENDIX B: McCrary and Balance Tests

In this section we carry out a McCrary test and conduct balance tests to investigate whether predetermined characteristics, such as genetic markers, are discontinuous around September 1, 1957.

**Appendix Figure B1:**
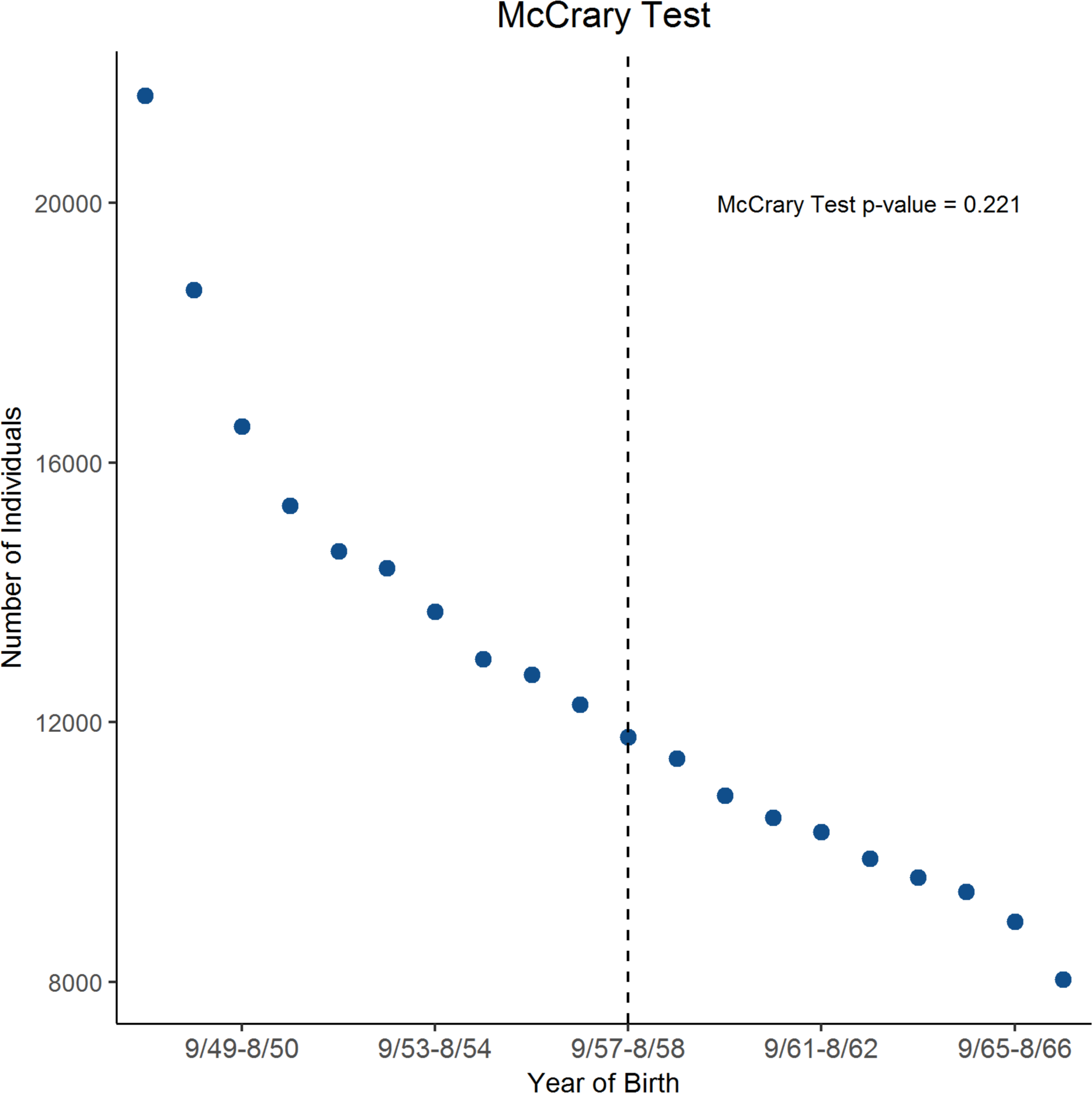
McCrary Test. The figure shows the fraction of study participants by year of birth. The dashed vertical line marks the first birth cohort affected by the 1972 school-leaving age reform. Cohorts born to the right of the line had to stay in school until age 16 while cohorts born before could leave at age 15. The estimated discontinuity of the density is −0.0206 with a standard error of 0.0169. *N* = 253,715.

**Appendix Figure B2:**
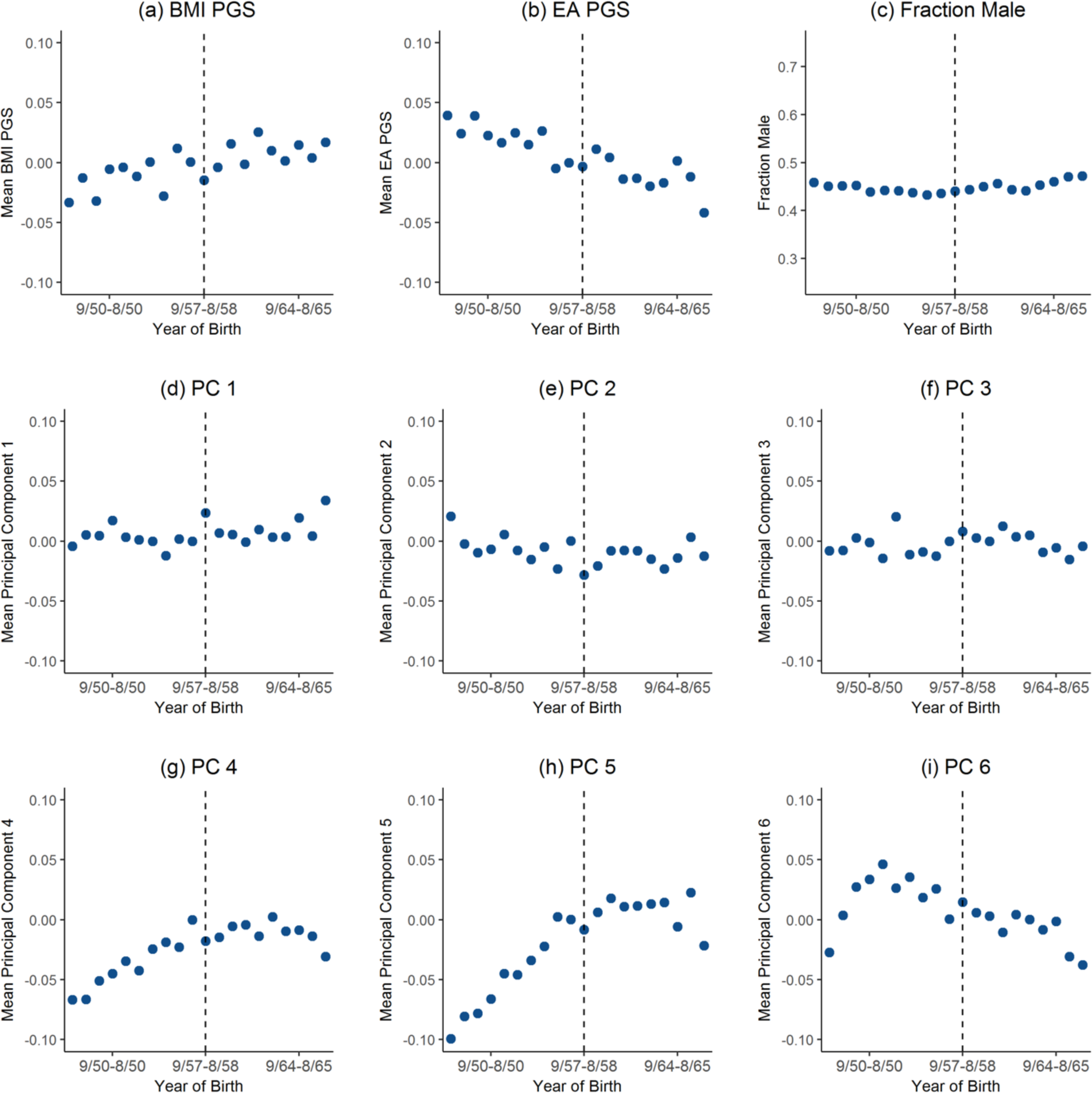
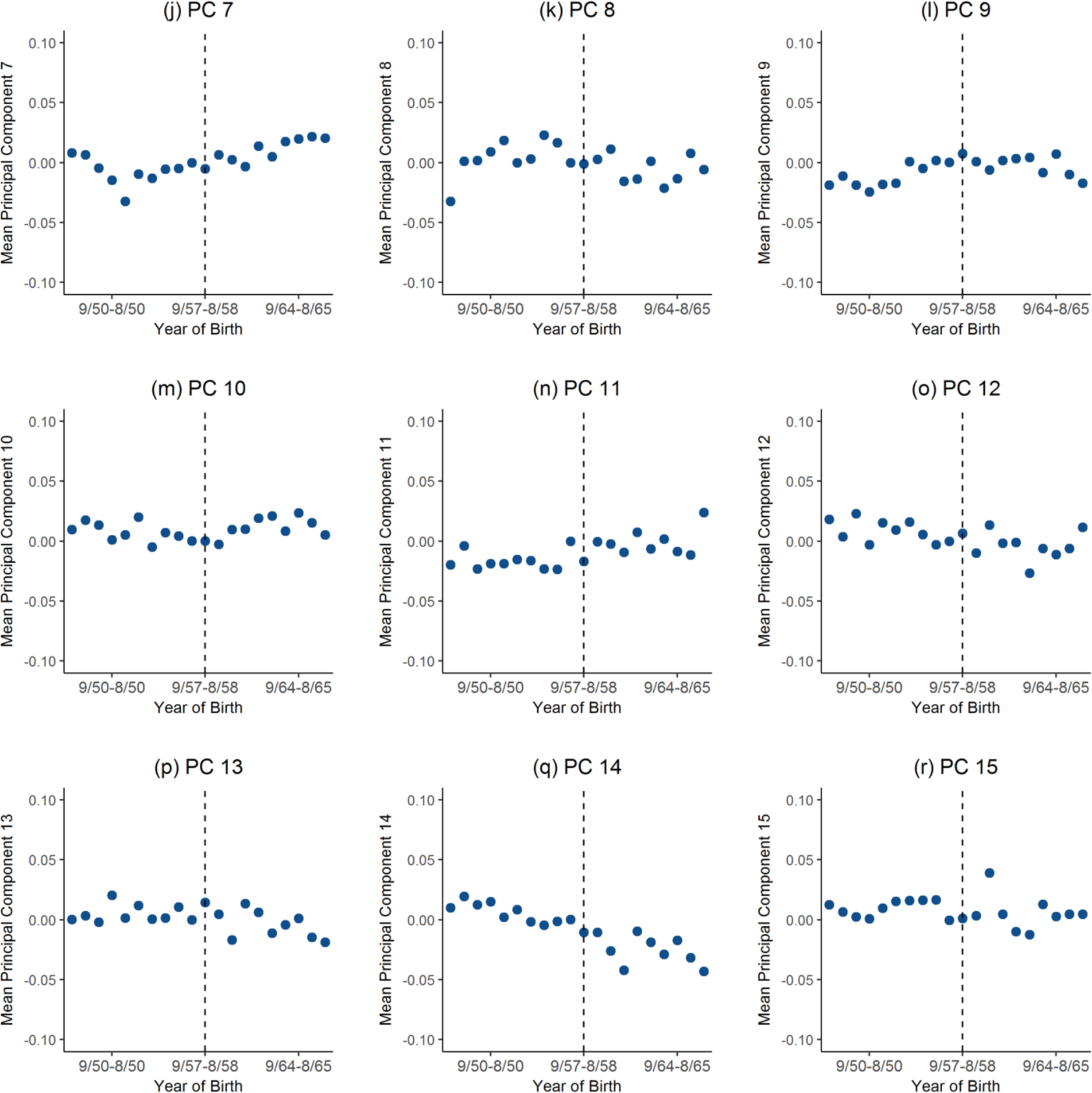
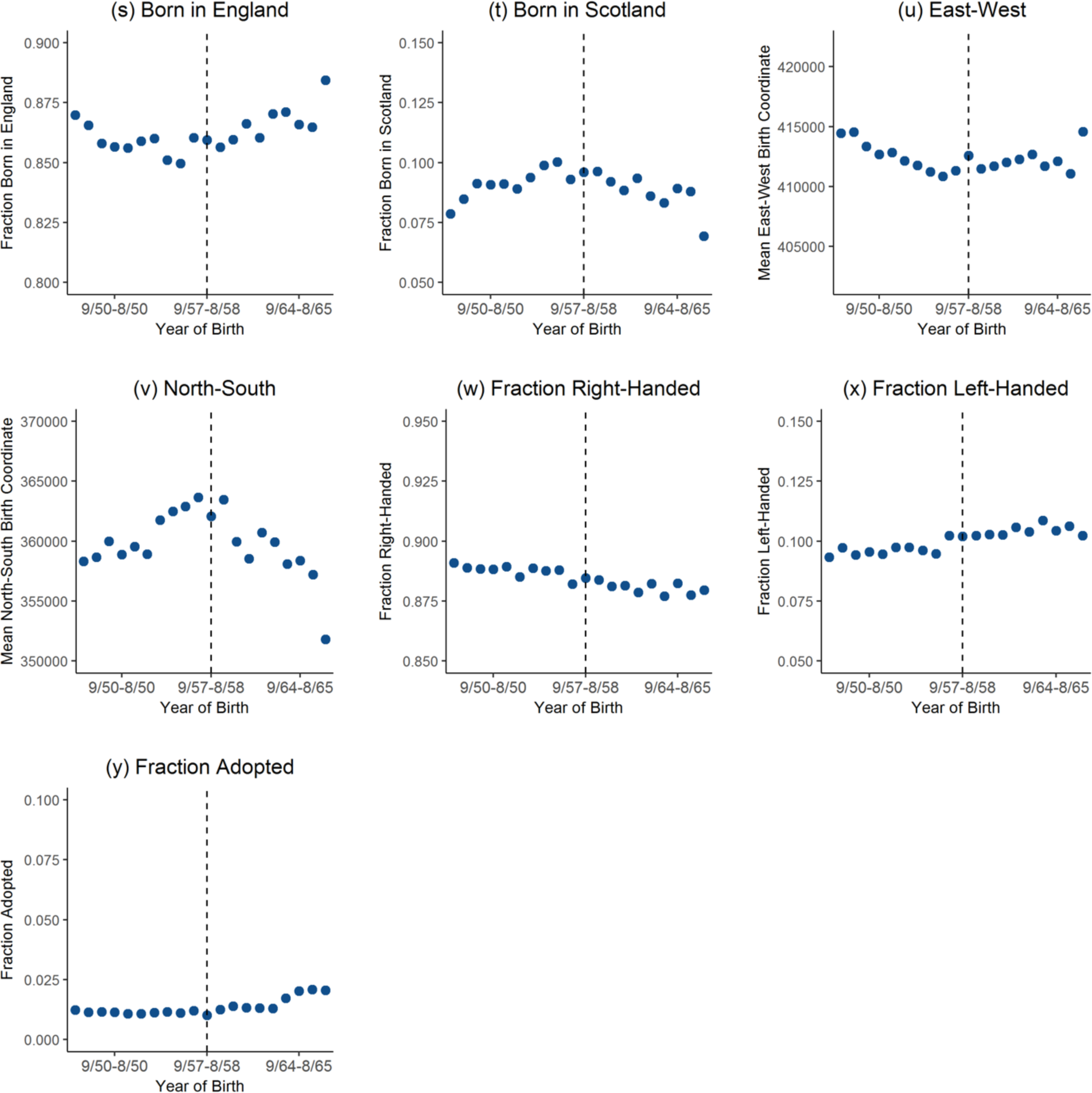
Balance Tests. The figures show averages by year of birth. The dashed vertical line marks the first birth cohort affected by the 1972 school-leaving age reform. Cohorts born to the right of the line had to stay in school until age 16 while cohorts born before could leave at age 15. PC1 to PC 15 refers to the first 15 principal components of the genotypic data. “East-West” and “North-South” correspond to the latitude and longitude coordinates of place of birth. *N* = 253,567 for all variables with the following exceptions: birthplace coordinates (*N* = 249,897); right- or left-handed (*N* = 253,519); and adopted (*N* = 253,279).

**Appendix Table B1:** Balance Tests. This table investigates whether predetermined characteristics are smooth around the September 1, 1957 cutoff. It reports the coefficient on an indicator for being born on or after September 1, 1957 (i.e., “Post”) from regressions where the dependent variables is listed in the column. The regressions include a quadratic polynomial in date of birth, which is allowed to be different before and after September 1, 1957. Robust standard errors. The mean of Y corresponds to the mean of the dependent variable among those born in the 12 months before September 1, 1957.

|  | *BMI*<br>*PGS* | *EA*<br>*PGS* | *Male* | *PC 1* | *PC 2* | *PC 3* | *PC 4* | *PC 5* | *PC 6* |
| --- | --- | --- | --- | --- | --- | --- | --- | --- | --- |
| Post | -0.02<br>[0.013] | 0.01<br>[0.013] | 0.01<br>[0.006] | 0.03<br>[0.013]** | -0.03<br>[0.013]** | 0.01<br>[0.013] | -0.02<br>[0.013] | -0.01<br>[0.013] | 0.02<br>[0.013]* |
| *N* | 253,567 | 253,567 | 253,567 | 253,567 | 253,567 | 253,567 | 253,567 | 253,567 | 253,567 |
| Mean of Y | 0.000 | 0.000 | 0.436 | 0.000 | 0.000 | 0.000 | 0.000 | 0.000 | 0.000 |
|  | *PC 7* | *PC 8* | *PC 9* | *PC 10* | *PC 11* | *PC 12* | *PC 13* | *PC 14* | *PC 15* |
| Post | -0.01<br>[0.013] | 0.00<br>[0.013] | 0.00<br>[0.013] | -0.01<br>[0.013] | -0.01<br>[0.013] | 0.01<br>[0.013] | 0.01<br>[0.013] | -0.01<br>[0.013] | 0.00<br>[0.013] |
| *N* | 253,567 | 253,567 | 253,567 | 253,567 | 253,567 | 253,567 | 253,567 | 253,567 | 253,567 |
| Mean of Y | 0.000 | 0.000 | 0.000 | 0.000 | 0.000 | 0.000 | 0.000 | 0.000 | 0.000 |
|  | *Born in*<br>*England* | *Born in*<br>*Scotland* | *East Coor.*<br>*Birthplace* | *North Coor.*<br>*Birthplace* | *Right-*<br>*Handed* | *Left-*<br>*Handed* | *Adopted* |  |  |
| Post | 0.00<br>[0.005] | 0.00<br>[0.004] | 977.14<br>[1,035.757] | -1348.92<br>[2,154.922] | 0.00<br>[0.004] | 0.00<br>[0.004] | 0.00<br>[0.001] |  |  |
| *N* | 253,567 | 253,567 | 249,897 | 249,897 | 253,519 | 253,519 | 253,279 |  |  |
| Mean of Y | 0.861 | 0.093 | 411,341 | 363,640 | 0.882 | 0.102 | 0.012 |  |  |

**The p-value for a joint test of the null hypothesis that there is no discontinuity for any of the 25 variables above is 0.6921.**

## APPENDIX C: Replication of Barcellos, Carvalho, and Turley (2017)

This section provides a summary of the results in Barcellos, Carvalho, and Turley (2017). Notice, however, that the results shown here are not identical to the results in Barcellos, Carvalho, and Turley (2017) because we use a slightly different sample. In particular, we drop from our sample non-whites and study participants with no genotypic data or with genotypic data that did not pass the quality controls.

**Appendix Table C1:** First Stage. This table estimates the effects of the 1972 school-leaving age reform on the fraction staying in school until age 16. The dependent variable is an indicator variable for whether participant stayed in school until at least age 16. “Post” in an indicator variable for being born on or after September 1, 1957. The regressions include a quadratic polynomial in date of birth, which is allowed to be different before and after September 1, 1957. The second column includes controls, namely male, age in days and age squared, dummies for calendar month of birth, and dummies for country of birth. Robust standard errors. *N* = 253,567.

|  | **1 if Stayed in School**<br>**until Age 16 or Older** |  |
| --- | --- | --- |
| Post | 0.151***<br>(0.004) | 0.140***<br>(0.004) |
| Controls? | N | Y |

**Appendix Figure C1:**
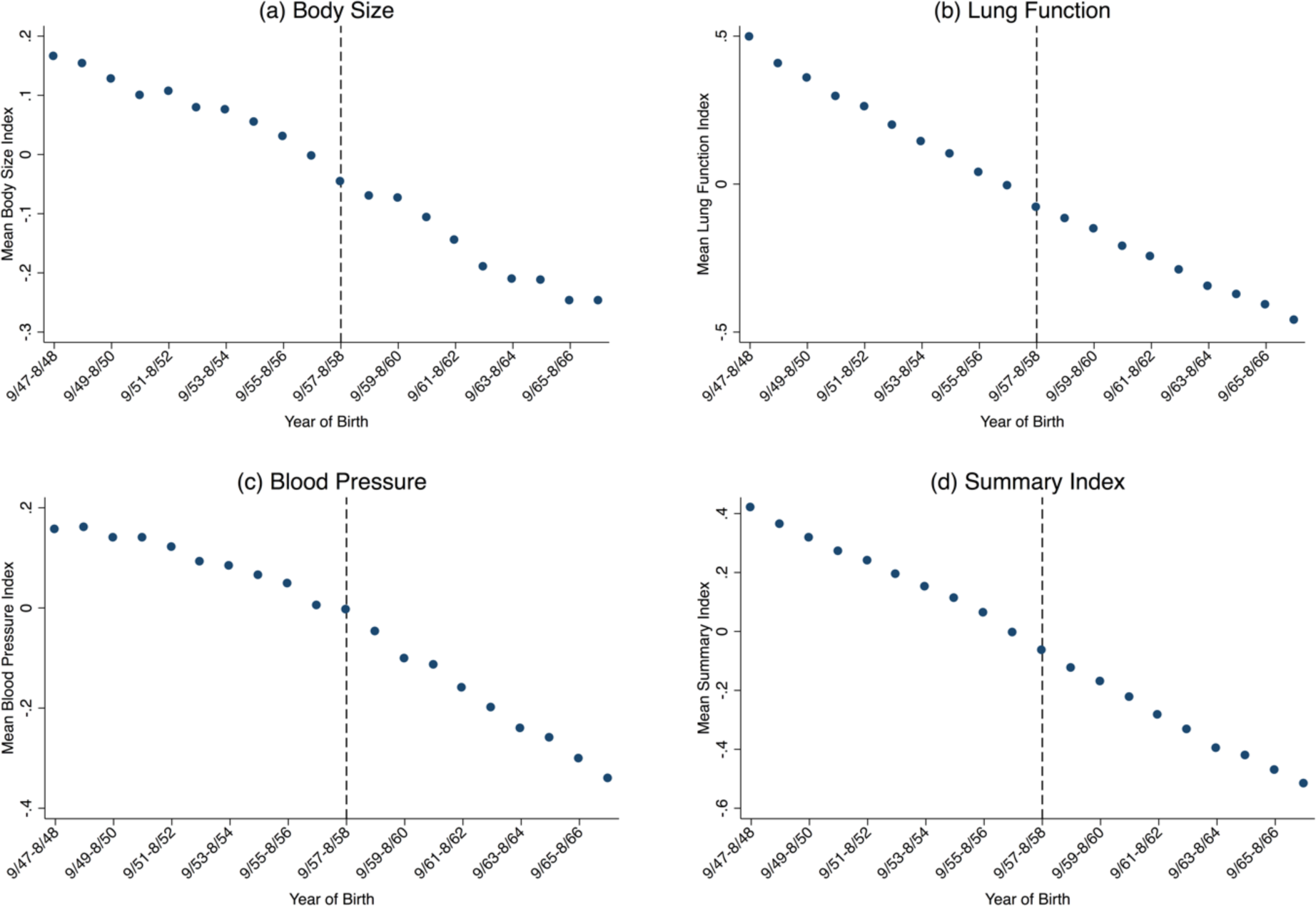
Mean Health Indices by Year Before and After the Reform’s Date-of-Birth Cutoff. The figures show averages of continuous measures of (a) body size (b) lung function (c) blood pressure and (d) summary indices by year of birth. The dashed vertical line marks the first birth cohort affected by the 1972 school-leaving age reform. Cohorts born to the right of the line had to stay in school until age 16 while cohorts born before could leave at age 15. *N* = 249,699 (a), 203,048 (b), 253,377 (c), 200,398 (d).

**Appendix Figure C2:**
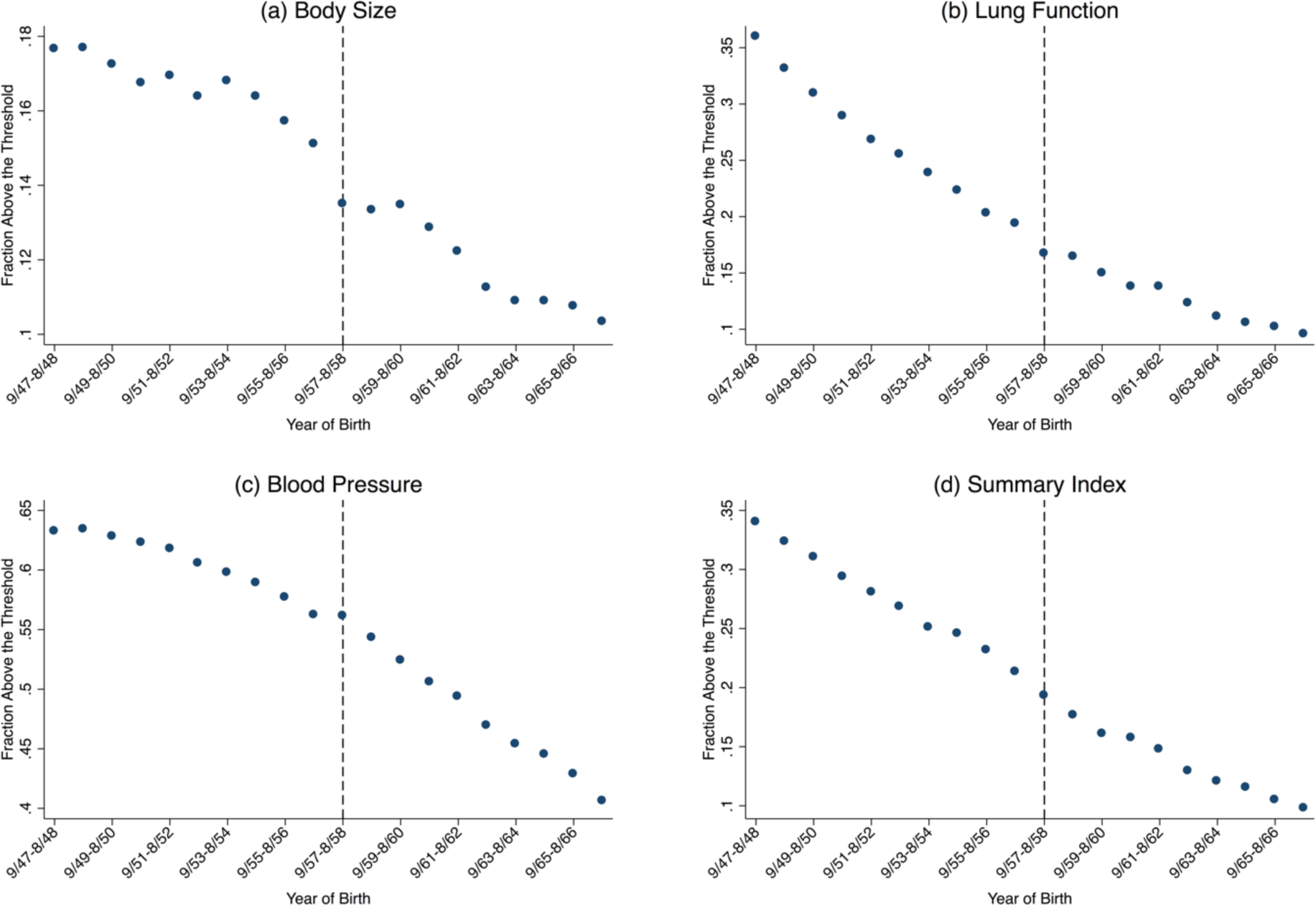
Fraction with a Health Index above the Pre-specified Threshold by Year Before and After the Reform’s Date-of-Birth Cutoff. The figures show averages of binary measures of (a) body size (b) lung function (c) blood pressure and (d) summary indices by year of birth. The dashed vertical line marks the first birth cohort affected by the 1972 school-leaving age reform. Cohorts born to the right of the line had to stay in school until age 16 while cohorts born before could leave at age 15. *N* = 249,699 (a), 203,048 (b), 253,377 (c), 200,398 (d).

**Appendix Table C2:** Effects of Education on Health. This table show estimates of the effects of education on health. The top panel shows reduced-form estimates of the effect of the 1972 raising of the school-leaving age reform. The bottom panel shows two-stage least squares (2SLS) estimates of the effect of staying in school until age 16. Above threshold is 1 if index is above the threshold specified in pre-analysis plan (see *Appendix* A). “Post” in an indicator variable for being born on or after September 1, 1957. Edu16 is an indicator for staying in school until age 16 and is instrumented by “Post”. The regressions include a quadratic polynomial in date of birth, which is allowed to be different before and after September 1, 1957, and controls for gender, age in days and age squared, dummies for calendar month of birth, and dummies for country of birth. Robust standard errors.

|  | Continuous |  |  |  | Above Threshold |  |  |  |
| --- | --- | --- | --- | --- | --- | --- | --- | --- |
|  | Body Size | Lung Function | Blood Pressure | Summary Index | Body Size | Lung Function | Blood Pressure | Summary Index |
| **Reduced-Form** |  |  |  |  |  |  |  |  |
| Post | -0.023*<br>(0.013) | -0.023<br>(0.015) | 0.016<br>(0.013) | -0.017<br>(0.015) | -0.012***<br>(0.005) | -0.013**<br>(0.006) | 0.014**<br>(0.006) | -0.013**<br>(0.006) |
| **Two-stage Least Squares** |  |  |  |  |  |  |  |  |
| Edu16 | -0.168*<br>(0.093) | -0.181<br>(0.114) | 0.118<br>(0.094) | -0.137<br>(0.114) | -0.087***<br>(0.033) | -0.102**<br>(0.044) | 0.103**<br>(0.047) | -0.099**<br>(0.046) |
| Observations | 249,699 | 203,048 | 253,377 | 200,398 | 249,699 | 203,048 | 253,377 | 200,398 |
| Dependent Variable Mean among Compliers | 0.261 | 0.269 | 0.090 | 0.311 | 0.215 | 0.287 | 0.611 | 0.317 |

## APPENDIX D: Mean Health Outcomes by the BMI and EA PGS

The figures in this section are comparable to Figure 1 in the body of the paper.

**Appendix Fig. D1.**
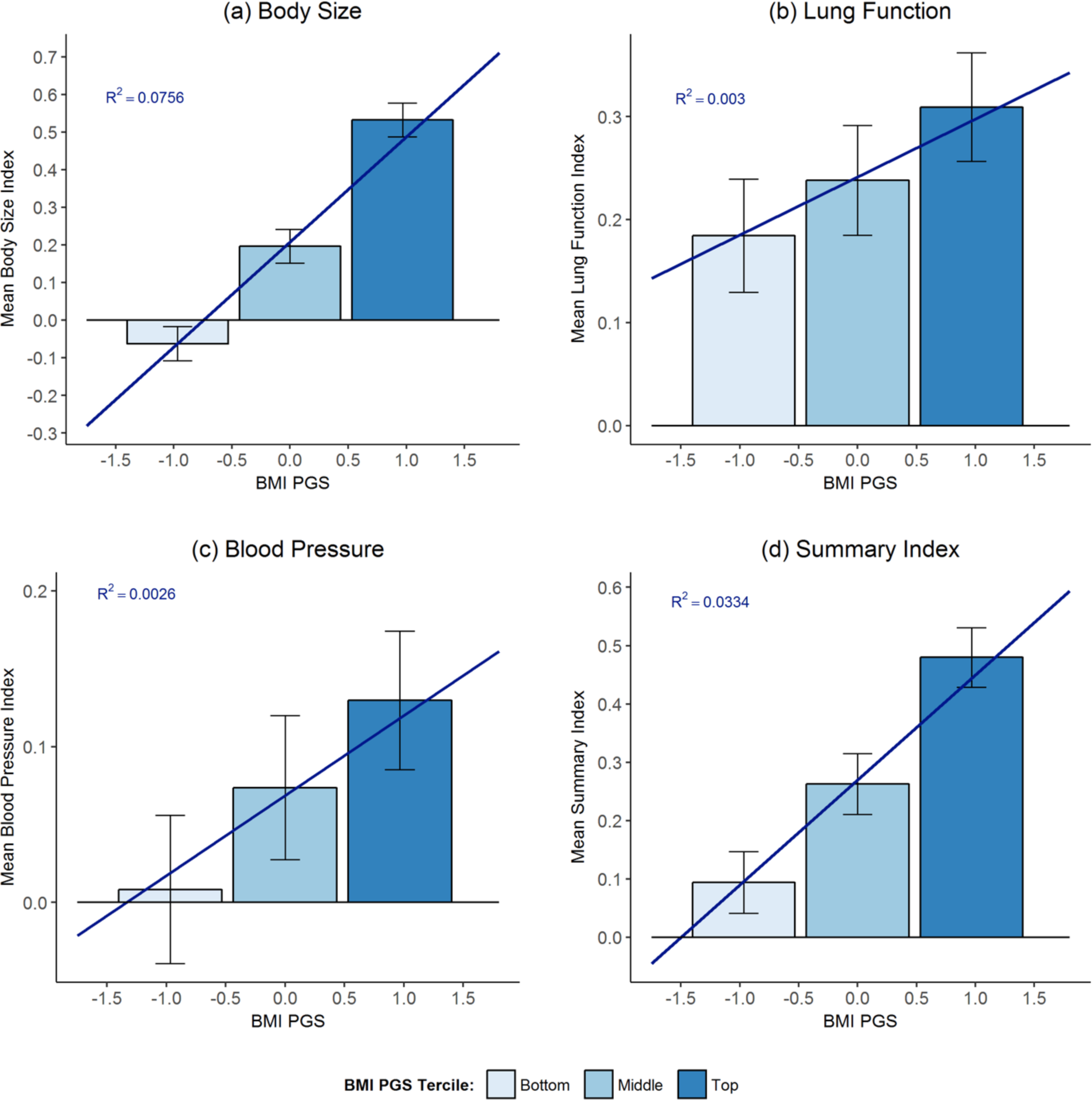
Mean Health Indices by BMI PGS. Bars show means of continuous measures of (**a**) body size, (**b**) lung function, (**c**) blood pressure, and (**d**) summary indices for the bottom, middle, and top terciles of the BMI PGS distribution with 95% confidence intervals. Sloped lines give linear projection of outcomes on BMI PGS. R^2^ gives the fraction of the variation in the outcome explained by the BMI PGS. We restrict the sample to participants born before September 1, 1957 who dropped out at age 15 or younger and control for a quadratic polynomial in date of birth.

**Appendix Fig. D2.**
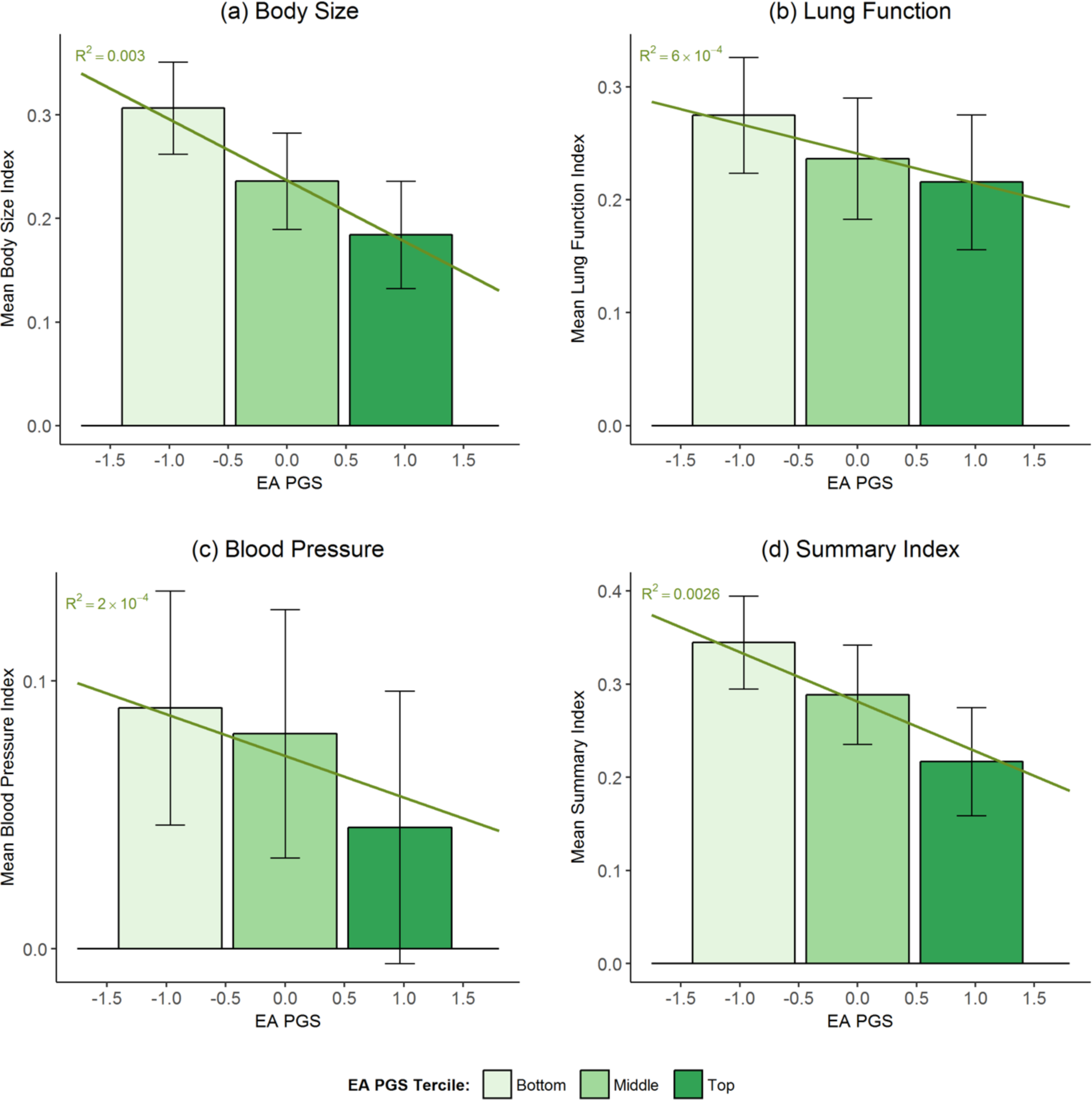
Mean Health Indices by BMI PGS. Bars show means of continuous measures of (**a**) body size, (**b**) lung function, (**c**) blood pressure, and (d) summary indices for the bottom, middle, and top terciles of the EA PGS distribution with 95% confidence intervals. Sloped lines give linear projection of outcomes on EA PGS. R^2^ gives the fraction of the variation in the outcome explained by the EA PGS. We restrict the sample to participants born before September 1, 1957 who dropped out at age 15 or younger and control for a quadratic polynomial in date of birth.

**Appendix Fig. D3.**
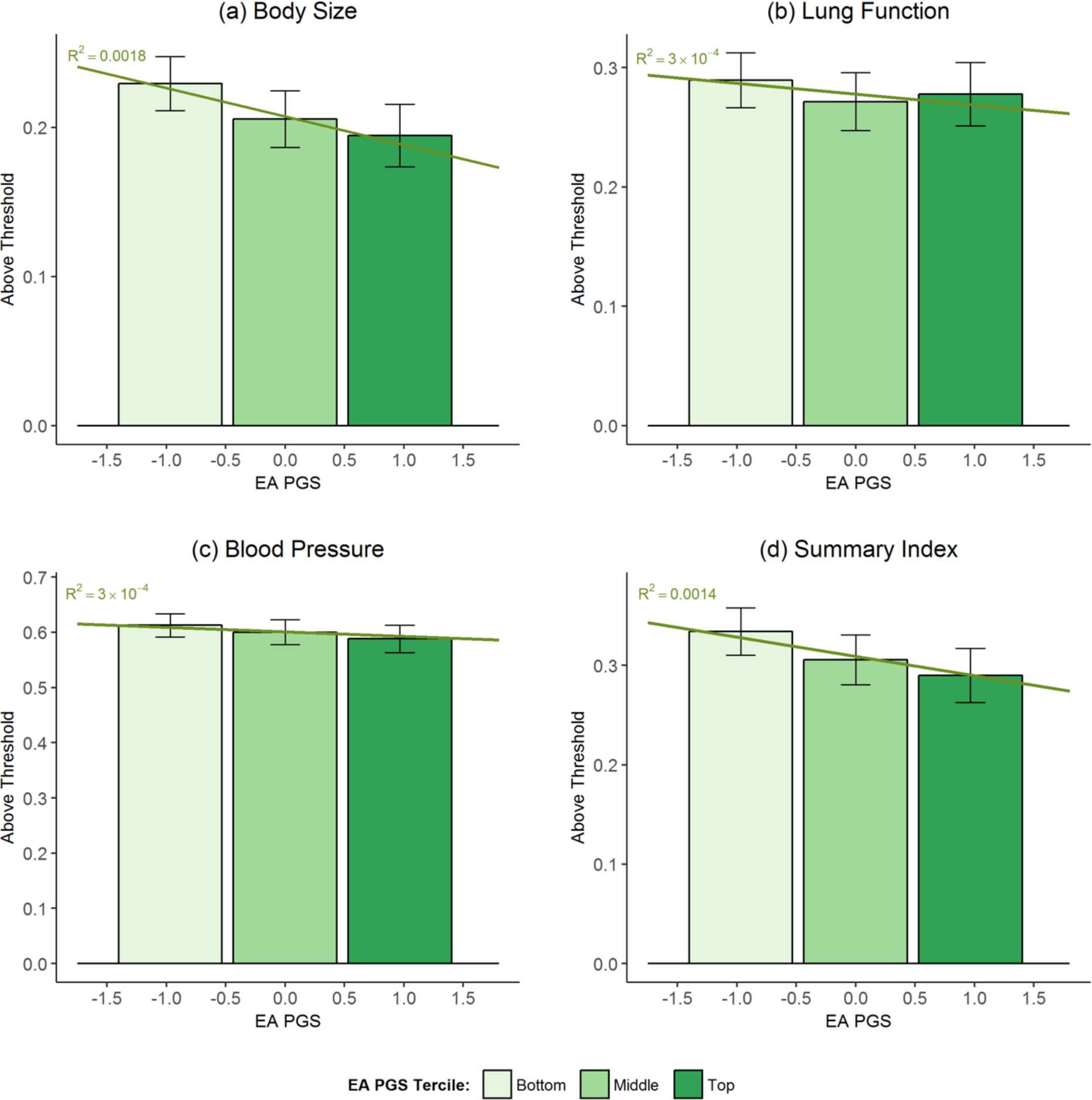
Fraction with Health Index above the Pre-specified threshold by EA PGS. Bars show means of binary measures of (**a**) body size, (**b**) lung function, (**c**) blood pressure, and (**d**) summary indices for the bottom, middle, and top terciles of the EA PGS distribution with 95% confidence intervals. Sloped lines give linear projection of outcomes on EA PGS. R^2^ gives the fraction of the variation in the outcome explained by the EA PGS. We restrict the sample to participants born before September 1, 1957 who dropped out at age 15 or younger and control for a quadratic polynomial in date of birth.

## APPENDIX E: First Stage and Reduced-Form Results

See Appendix Table C1 for first stage results that do not break by genetic makeup.

**Appendix Table E1:**
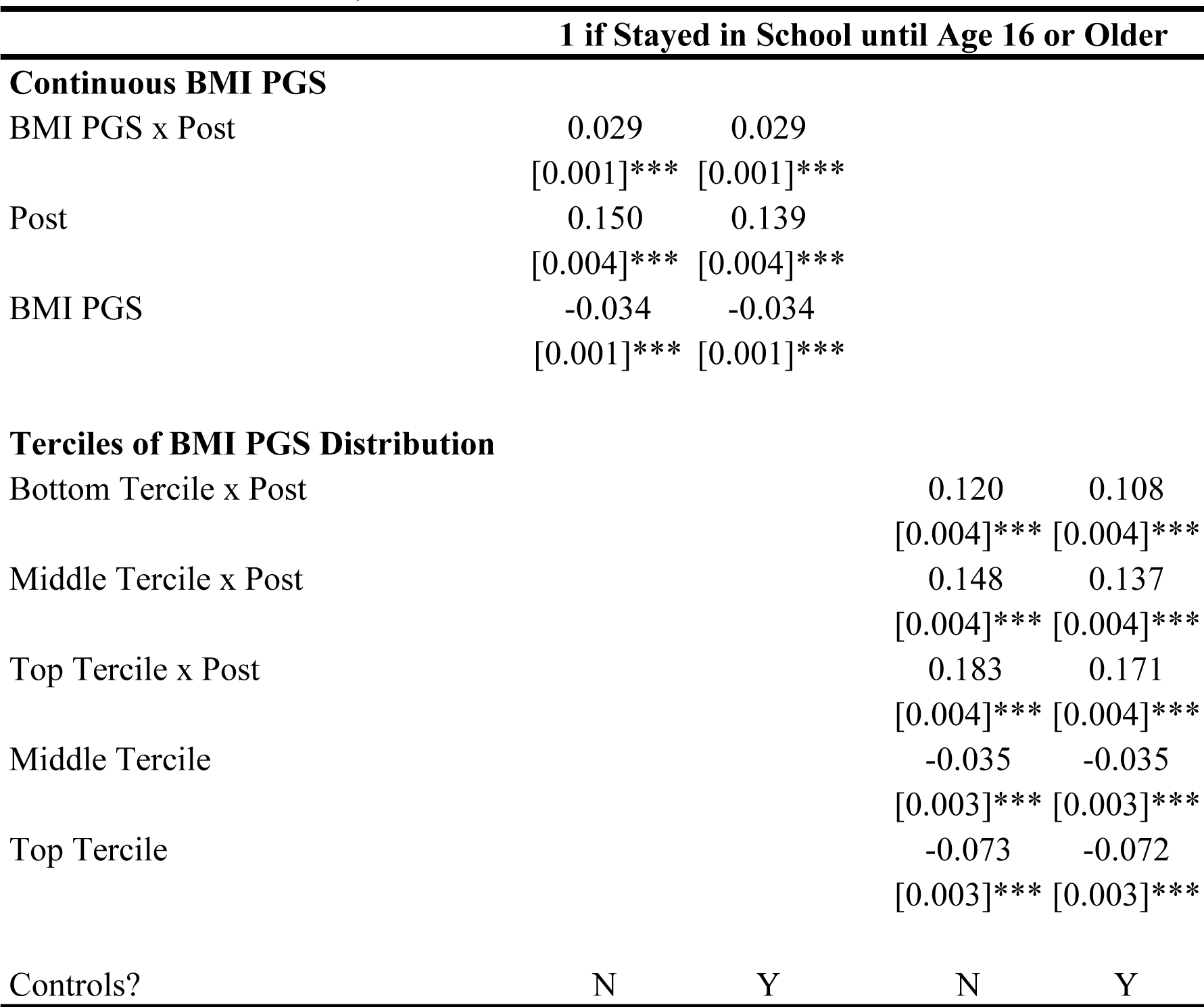
Does the Effect of the 1972 Raising of the School Leaving-Age on Education depend on the BMI PGS? This table investigates whether the effects of the 1972 school-leaving age reform on the fraction staying in school until age 16 depend on the BMI PGS. The dependent variable is an indicator variable for whether participant stayed in school until at least age 16. “Post” in an indicator variable for being born on or after September 1, 1957. “Bottom Tercile”, “Middle Tercile”, and “Top Tercile” are indicator variables for whether participant was in the bottom, middle, or top tercile of the BMI PGS distribution. The regressions include a quadratic polynomial in date of birth, which is allowed to be different before and after September 1, 1957; the first 15 principal components of the genotypic data; and the interactions of those principal components with “Post.” Even columns include controls, namely male, age in days and age squared, dummies for calendar month of birth, and dummies for country of birth. Robust standard errors. *N* = 253,567.

**Appendix Table E2:**
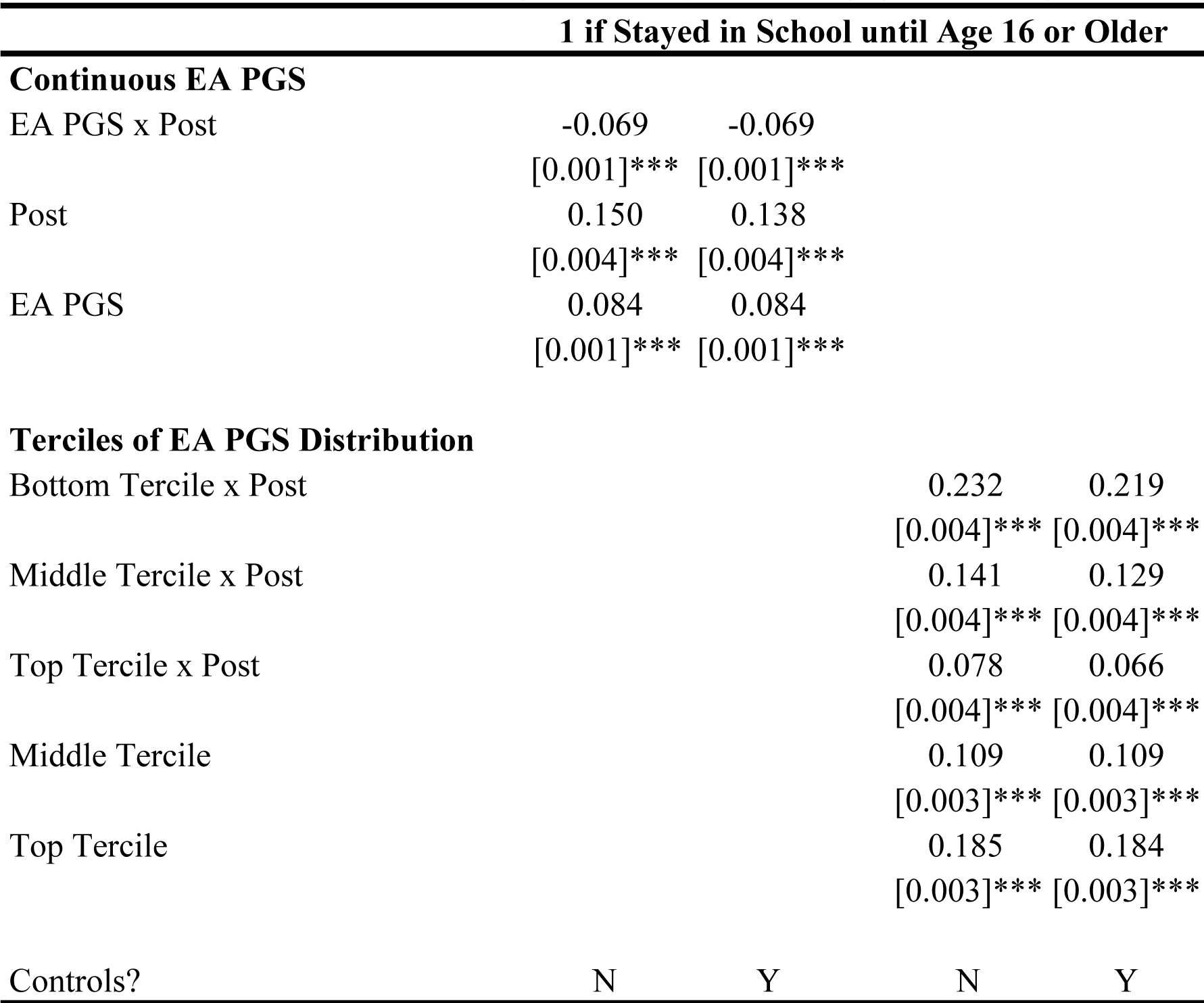
Does the Effect of the 1972 Raising of the School Leaving-Age on Education depend on the EA PGS? This table investigates whether the effects of the 1972 school-leaving age reform on the fraction staying in school until age 16 depend on the EA PGS. The dependent variable is an indicator variable for whether participant stayed in school until at least age 16. “Post” in an indicator variable for being born on or after September 1, 1957. “Bottom Tercile”, “Middle Tercile”, and “Top Tercile” are indicator variables for whether participant was in the bottom, middle, or top tercile of the EA PGS distribution. The regressions include a quadratic polynomial in date of birth, which is allowed to be different before and after September 1, 1957; the first 15 principal components of the genotypic data; and the interactions of those principal components with “Post.” Even columns include controls, namely male, age in days and age squared, dummies for calendar month of birth, and dummies for country of birth. Robust standard errors. *N* = 253,567.

**Appendix Table E3:** Effects of the 1972 Raising of the School-Leaving Age Reform on Health Reduced-. form estimates. Above threshold is 1 if index is above the threshold specified in pre-analysis plan (*see* *Appendix* A). “Post” in an indicator variable for being born on or after September 1, 1957. The regressions include a quadratic polynomial in date of birth, which is allowed to be different before and after September 1, 1957; the first 15 principal components of the genotypic data; the interactions of those principal components with “Post”; and controls for age, age in days and age squared, dummies for calendar month of birth, and dummies for country of birth. Robust standard errors. The “P-value for H_0_: No Effect of Education” is the p-value from a joint test that β_1_*= β_2_* = 0. The last row shows means of the dependent variable among pre-reform compliers, defined as individuals born before September 1, 1957 who dropped out before age 16.

|  | Continuous |  |  |  | Above Threshold |  |  |  |
| --- | --- | --- | --- | --- | --- | --- | --- | --- |
|  | Body Size | Lung Function | Blood Pressure | Summary Index | Body Size | Lung Function | Blood Pressure | Summary Index |
| **Interaction with BMI PGS** |  |  |  |  |  |  |  |  |
| BMI PGS × Post | 0.001<br>(0.005) | -0.018***<br>(0.005) | 0.015***<br>(0.005) | -0.004<br>(0.005) | -0.011***<br>(0.002) | -0.008***<br>(0.002) | 0.006**<br>(0.002) | -0.017***<br>(0.002) |
| Post | -0.015<br>(0.012) | -0.020<br>(0.015) | 0.018<br>(0.013) | -0.011<br>(0.014) | -0.010**<br>(0.005) | -0.012**<br>(0.006) | 0.015**<br>(0.006) | -0.011*<br>(0.006) |
| BMI PGS | 0.290***<br>(0.003) | 0.057***<br>(0.003) | 0.079***<br>(0.003) | 0.200***<br>(0.003) | 0.080***<br>(0.001) | 0.020***<br>(0.001) | 0.034***<br>(0.001) | 0.070***<br>(0.001) |
| P-value for $H_0$: No Effect of Education | 0.431 | 0.002 | 0.002 | 0.554 | $6.75 \times 10^{-10}$ | $4.69 \times 10^{-5}$ | 0.003 | $8.32 \times 10^{-15}$ |
| **Interaction with EA PGS** |  |  |  |  |  |  |  |  |
| EA PGS × Post | -0.002<br>(0.005) | 0.011**<br>(0.005) | -0.012***<br>(0.005) | 0.001<br>(0.005) | 0.004**<br>(0.002) | 0.010***<br>(0.002) | -0.005*<br>(0.002) | 0.012***<br>(0.002) |
| Post | -0.021<br>(0.013) | -0.021<br>(0.015) | 0.017<br>(0.013) | -0.015<br>(0.014) | -0.011**<br>(0.005) | -0.012**<br>(0.006) | 0.015**<br>(0.006) | -0.012**<br>(0.006) |
| EA PGS | -0.126***<br>(0.003) | -0.074***<br>(0.003) | -0.063***<br>(0.003) | -0.131***<br>(0.003) | -0.031***<br>(0.001) | -0.025***<br>(0.001) | -0.027***<br>(0.002) | -0.043***<br>(0.001) |
| P-value for $H_0$: No Effect of Education | 0.241 | 0.043 | 0.015 | 0.587 | 0.004 | $4.90 \times 10^{-7}$ | 0.011 | $1.12 \times 10^{-8}$ |
| Observations | 249,699 | 203,048 | 253,377 | 200,398 | 249,699 | 203,048 | 253,377 | 200,398 |
| Dependent Variable Mean among Compliers | 0.261 | 0.269 | 0.090 | 0.311 | 0.215 | 0.287 | 0.611 | 0.317 |

Appendix Table E3 corresponds to the reduced-form version of Table 1 shown in the body of the paper.

## APPENDIX F: The Two-Stage Least Squares Effect of an Additional Year of Schooling at Age 16 on Health Indices by the BMI and EA PGS

**Appendix Figure F1.**
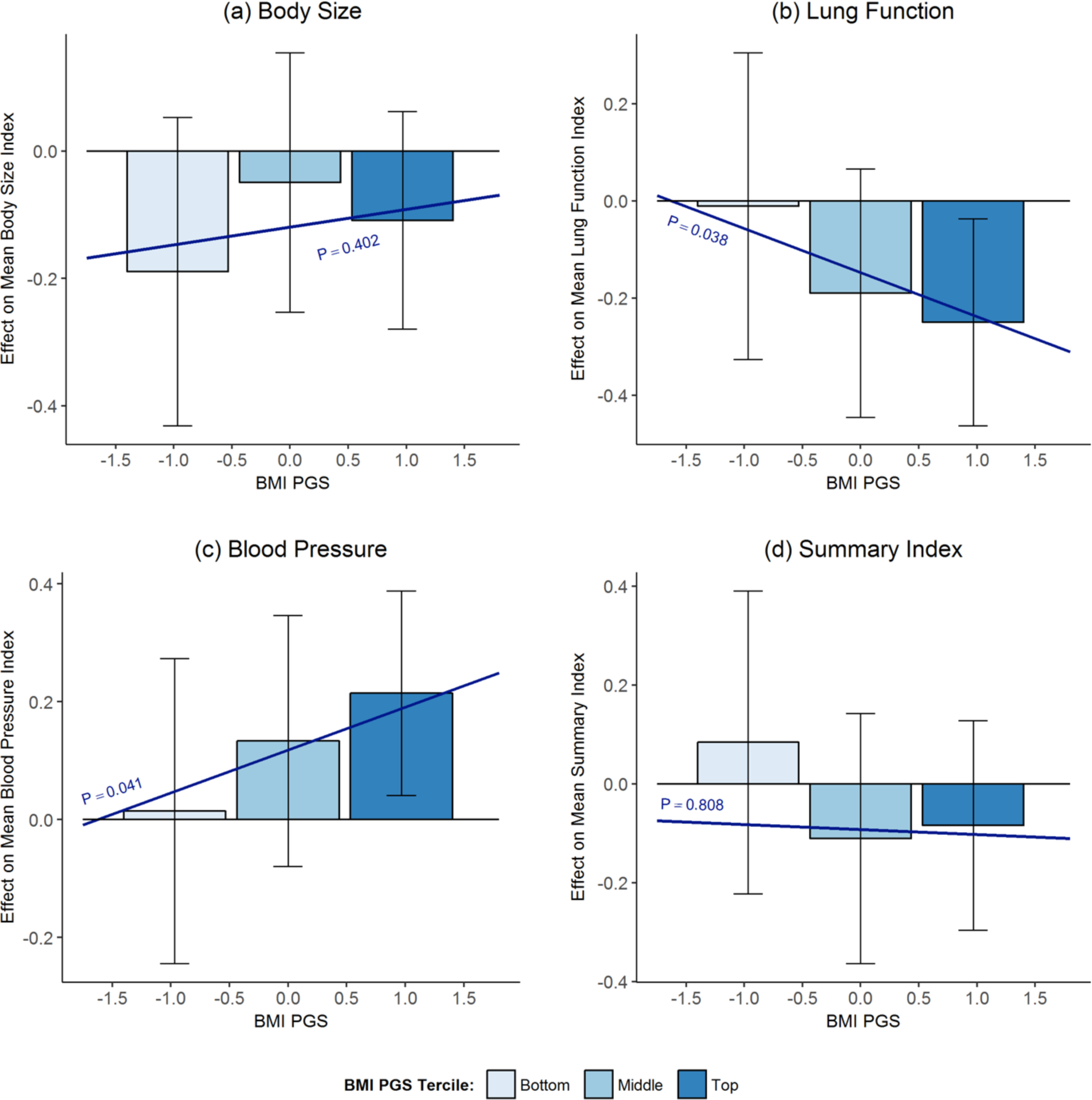
The 2SLS Effect of an Additional Year of Schooling at Age 16 on Health Indices by BMI PGS. Bars show 2SLS point estimates of effect of staying in school until age 16 on continuous measures of (**a**) body size (**b**) lung function (**c**) blood pressure and (d) summary indices for the bottom, middle, and top terciles of the BMI PGS distribution. Brackets show 95% confidence intervals. Sloped lines plot 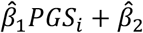. “P” corresponds to the p-value of *H_0:_ β_1_* = 0. The regressions include a quadratic polynomial in date of birth, which is allowed to be different before and after September 1, 1957; the first 15 principal components of the genotypic data; the interactions of those principal components with “Edu 16”; and controls for age, age in days and age squared, dummies for calendar month of birth, and dummies for country of birth.

**Appendix Figure F2.**
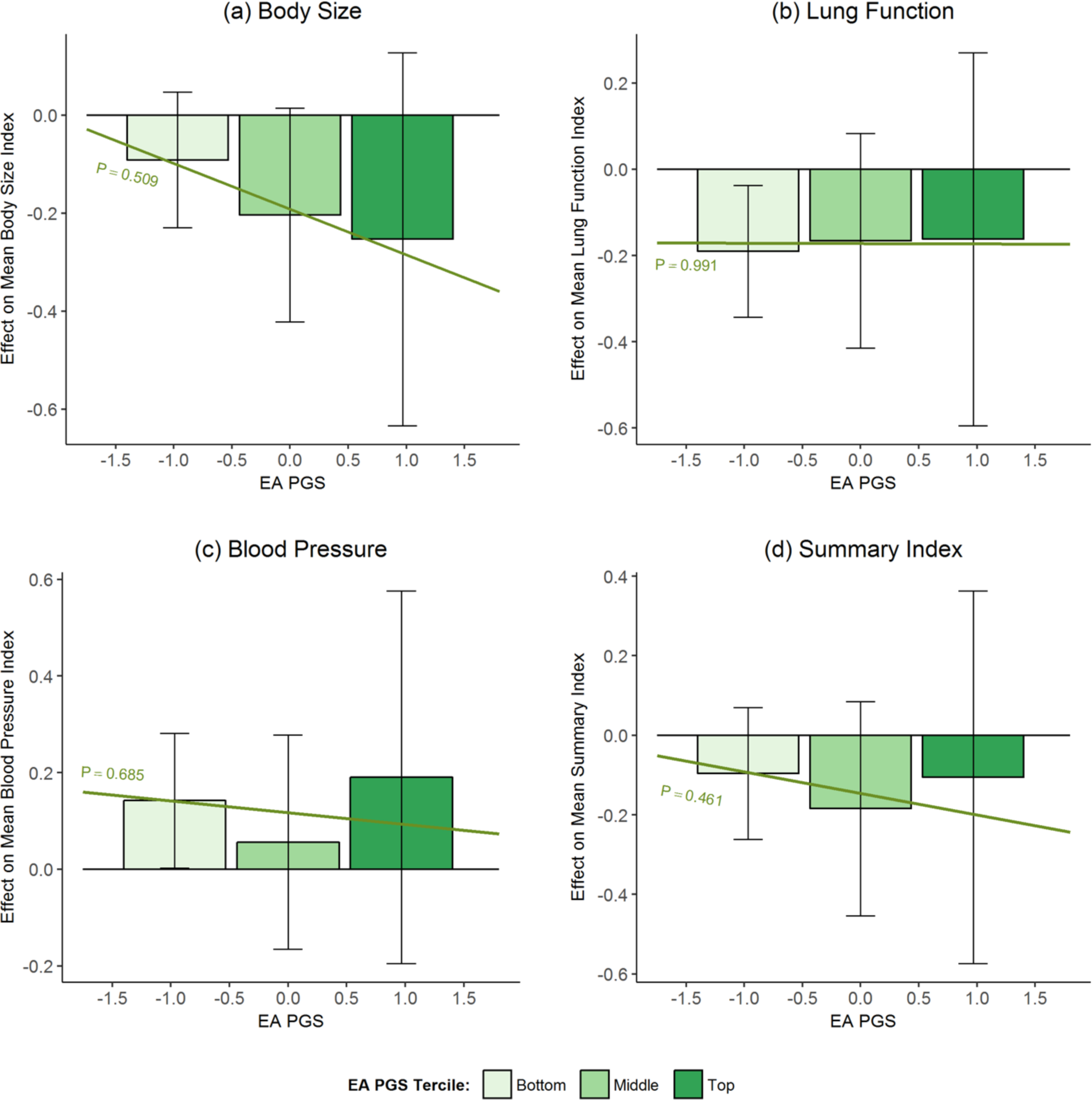
The 2SLS Effect of an Additional Year of Schooling at Age 16 on Health Indices by EA PGS. Bars show 2SLS point estimates of effect of staying in school until age 16 on continuous measures of (**a**) body size (**b**) lung function (**c**) blood pressure and (**d**) summary indices for the bottom, middle, and top terciles of the EA PGS distribution. Brackets show 95% confidence intervals. Sloped lines plot 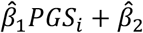. “P” corresponds to the p-value of *H_0_:β_1_* = 0. The regressions include a quadratic polynomial in date of birth, which is allowed to be different before and after September 1, 1957; the first 15 principal components of the genotypic data; the interactions of those principal components with “Edu 16”; and controls for age, age in days and age squared, dummies for calendar month of birth, and dummies for country of birth.

**Appendix Figure F3.**
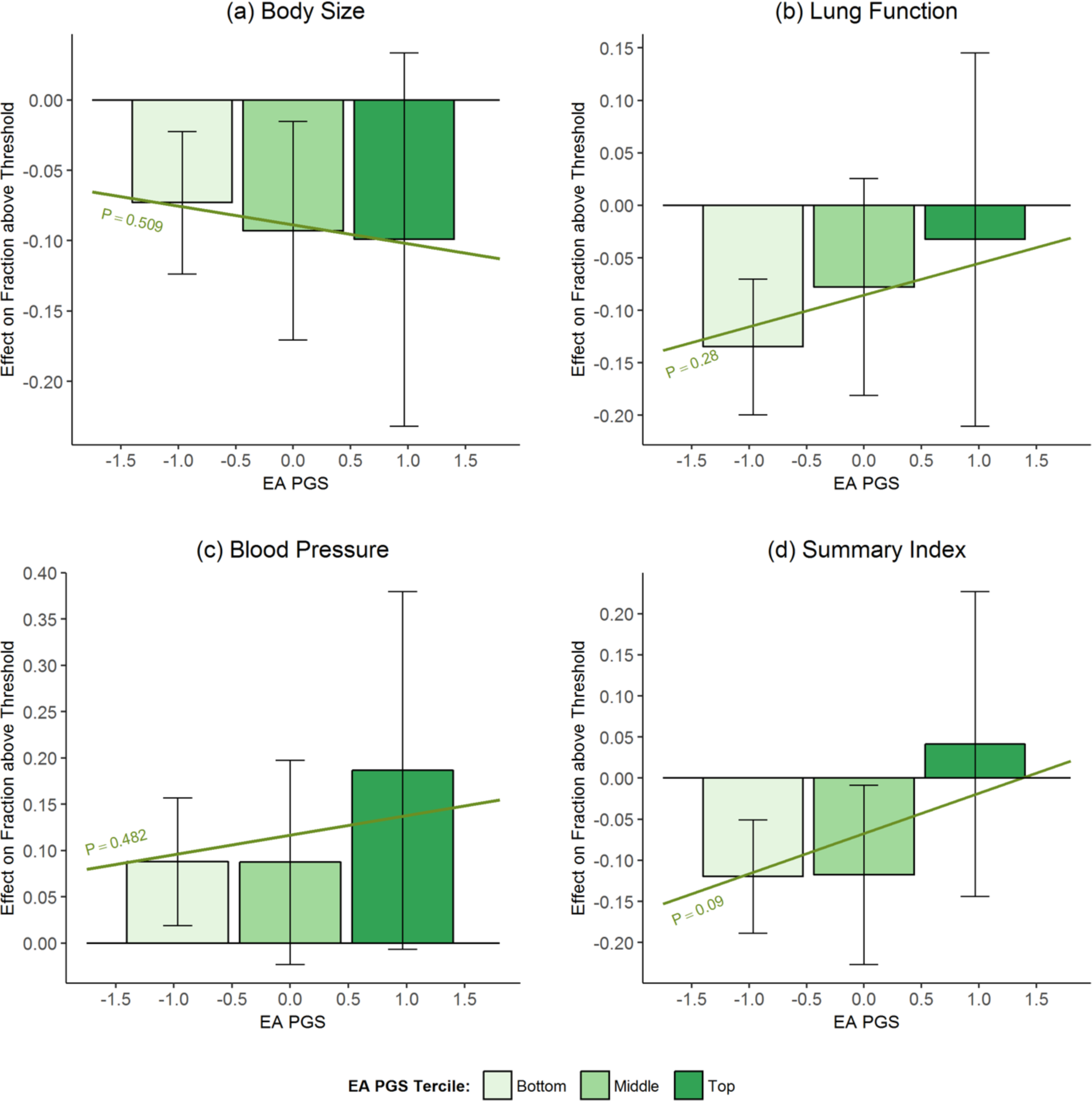
The 2SLS Effect of an Additional Year of Schooling at Age 16 on the Fraction with a Health Index above the Pre-specified Threshold by BMI PGS. Bars show 2SLS point estimates of effect of staying in school until age 16 on binary measures of (**a**) body size (**b**) lung function (**c**) blood pressure and (**d**) summary indices for the bottom, middle, and top terciles of the EA PGS distribution. Brackets show 95% confidence intervals. Sloped lines plot 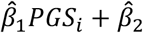. “P” corresponds to the p-value of *H_0:_ β_1_* = 0. The regressions include a quadratic polynomial in date of birth, which is allowed to be different before and after September 1, 1957; the first 15 principal components of the genotypic data; the interactions of those principal components with “Edu 16”; and controls for age, age in days and age squared, dummies for calendar month of birth, and dummies for country of birth.

## APPENDIX G: Sensitivity to Simultaneous Inclusion of BMI and EA PGSs and to Exclusion of Controls

**Appendix Table G1.** Sensitivity to Simultaneous Inclusion of BMI and EA PGSs. 2SLS estimates. Above threshold is 1 if index is above the threshold specified in pre-analysis plan (*see* *Appendix* A). Edu16 is an indicator for staying in school until age 16 and is instrumented by an indicator for being born after September 1, 1957. The regressions include a quadratic polynomial in date of birth, which is allowed to be different before and after September 1, 1957; the first 15 principal components of the genotypic data; the interactions of those principal components with “Edu 16”; and controls for age, age in days and age squared, dummies for calendar month of birth, and dummies for country of birth. Robust standard errors. The “P-value for H_0_: No Effect of Education” is the p-value from a joint test that the coefficients on “BMI PGS x Edu16”, “EA PGS x Edu16 “, and on “Edu16” are all equal to zero. The last row shows means of the dependent variable among pre-reform compliers, defined as individuals born before September 1, 1957 who dropped out before age 16.

|  | Continuous |  |  |  | Above Threshold |  |  |  |
| --- | --- | --- | --- | --- | --- | --- | --- | --- |
|  | Body Size | Lung Function | Blood Pressure | Summary Index | Body Size | Lung Function | Blood Pressure | Summary Index |
| BMI PGS × Edu16 | 0.014<br>[0.031] | -0.095<br>[0.039]** | 0.070<br>[0.032]** | -0.020<br>[0.038] | -0.060<br>[0.011]*** | -0.033<br>[0.015]** | 0.021<br>[0.016] | -0.095<br>[0.015]*** |
| EA PGS × Edu16 | -0.065<br>[0.056] | -0.016<br>[0.073] | -0.004<br>[0.059] | -0.040<br>[0.071] | -0.019<br>[0.020] | 0.025<br>[0.027] | 0.028<br>[0.029] | 0.036<br>[0.028] |
| Edu16 | -0.143<br>[0.115] | -0.153<br>[0.149] | 0.120<br>[0.122] | -0.107<br>[0.146] | -0.068<br>[0.042] | -0.079<br>[0.057] | 0.119<br>[0.061]* | -0.043<br>[0.059] |
| BMI PGS | 0.262<br>[0.027]*** | 0.118<br>[0.035]*** | 0.013<br>[0.029] | 0.195<br>[0.034]*** | 0.125<br>[0.010]*** | 0.041<br>[0.014]*** | 0.015<br>[0.014] | 0.141<br>[0.014]*** |
| EA PGS | 0.003<br>[0.054] | -0.039<br>[0.071] | -0.054<br>[0.057] | -0.049<br>[0.069] | 0.008<br>[0.019] | -0.035<br>[0.027] | -0.052<br>[0.028]* | -0.054<br>[0.028]* |
| P-value for $H_0$: No Effect of Education | 0.657 | 0.002 | 0.001 | 0.722 | $2.52 \times 10^{-9}$ | $3.78 \times 10^{-8}$ | 0.005 | $1.34 \times 10^{-17}$ |
| Observations | 249,699 | 203,048 | 253,377 | 200,398 | 249,699 | 203,048 | 253,377 | 200,398 |
| Dependent Variable Mean among Compliers | 0.261 | 0.269 | 0.090 | 0.311 | 0.215 | 0.287 | 0.611 | 0.317 |

Appendix Table G1 corresponds to the results shown in Table 1 in the body of the paper when both the BMI and EA PGSs are included in the regression - see section VII. Genetic Heterogeneity in Appendix A.

**Appendix Table G2.** Sensitivity to Exclusion of Controls. 2SLS estimates. Above threshold is 1 if index is above the threshold specified in pre-analysis plan (see *Appendix* A). Edu16 is an indicator for staying in school until age 16 and is instrumented by an indicator for being born after September 1, 1957. The regressions include a quadratic polynomial in date of birth, which is allowed to be different before and after September 1, 1957; the first 15 principal components of the genotypic data; and the interactions of those principal components with “Edu 16”. It excludes controls for age, age in days and age squared, dummies for calendar month of birth, and dummies for country of birth. Robust standard errors. The “P-value for H_0_: No Effect of Education” is the p-value from a joint test that β_1_ = β_2_= 0. The last row shows means of the dependent variable among pre-reform compliers, defined as individuals born before September 1, 1957 who dropped out before age 16.

|  | Continuous |  |  |  | Above Threshold |  |  |  |
| --- | --- | --- | --- | --- | --- | --- | --- | --- |
|  | Body Size | Lung Function | Blood Pressure | Summary Index | Body Size | Lung Function | Blood Pressure | Summary Index |
| **Interaction with BMI PGS** |  |  |  |  |  |  |  |  |
| BMI PGS × Edu16 | 0.029<br>(0.033) | -0.089**<br>(0.043) | 0.072**<br>(0.035) | -0.009<br>(0.041) | -0.057***<br>(0.011) | -0.037**<br>(0.016) | 0.017<br>(0.017) | -0.101***<br>(0.016) |
| Edu16 | -0.121<br>(0.088) | -0.153<br>(0.113) | 0.121<br>(0.094) | -0.096<br>(0.111) | -0.056*<br>(0.032) | -0.088**<br>(0.043) | 0.105**<br>(0.047) | -0.059<br>(0.045) |
| BMI PGS | 0.263***<br>(0.030) | 0.125***<br>(0.039) | 0.025<br>(0.032) | 0.205***<br>(0.037) | 0.125***<br>(0.010) | 0.048***<br>(0.015) | 0.025<br>(0.016) | 0.151***<br>(0.015) |
| P-value for $H_0$: No Effect of Education | 0.382 | 0.001 | 0.002 | 0.509 | $8.55 \times 10^{-10}$ | $4.46 \times 10^{-5}$ | 0.002 | $1.55 \times 10^{-14}$ |
| **Interaction with EA PGS** |  |  |  |  |  |  |  |  |
| EA PGS × Edu16 | -0.095*<br>(0.055) | -0.006<br>(0.069) | -0.023<br>(0.056) | -0.058<br>(0.069) | -0.011<br>(0.019) | 0.030<br>(0.026) | 0.020<br>(0.028) | 0.048*<br>(0.027) |
| Edu16 | -0.191*<br>(0.109) | -0.179<br>(0.134) | 0.120<br>(0.111) | -0.148<br>(0.133) | -0.082**<br>(0.038) | -0.085*<br>(0.051) | 0.114**<br>(0.055) | -0.068<br>(0.053) |
| EA PGS | -0.035<br>(0.053) | -0.054<br>(0.067) | -0.055<br>(0.054) | -0.073<br>(0.067) | -0.015<br>(0.019) | -0.042*<br>(0.025) | -0.053*<br>(0.027) | -0.077***<br>(0.026) |
| P-value for $H_0$: No Effect of Education | 0.198 | 0.032 | 0.013 | 0.517 | 0.007 | $2.85 \times 10^{-7}$ | 0.009 | $2.16 \times 10^{-8}$ |
| Observations | 249,699 | 203,048 | 253,377 | 200,398 | 249,699 | 203,048 | 253,377 | 200,398 |
| Dependent Variable Mean among Compliers | 0.261 | 0.269 | 0.090 | 0.311 | 0.215 | 0.287 | 0.611 | 0.317 |

Appendix Table G2 corresponds to Table 1 in the body of the paper when we drop controls for gender, age in days, age squared, dummies for calendar month of birth, and dummies for country of birth.

## APPENDIX H: Robustness to Different Bandwidths and Linear Trends

The figures in this section investigate the sensitivity of our results to using linear trends instead of quadratic trends and to using shorter bandwidths of 3 or 5 years instead of 10 years.

**Appendix Fig. H1.**
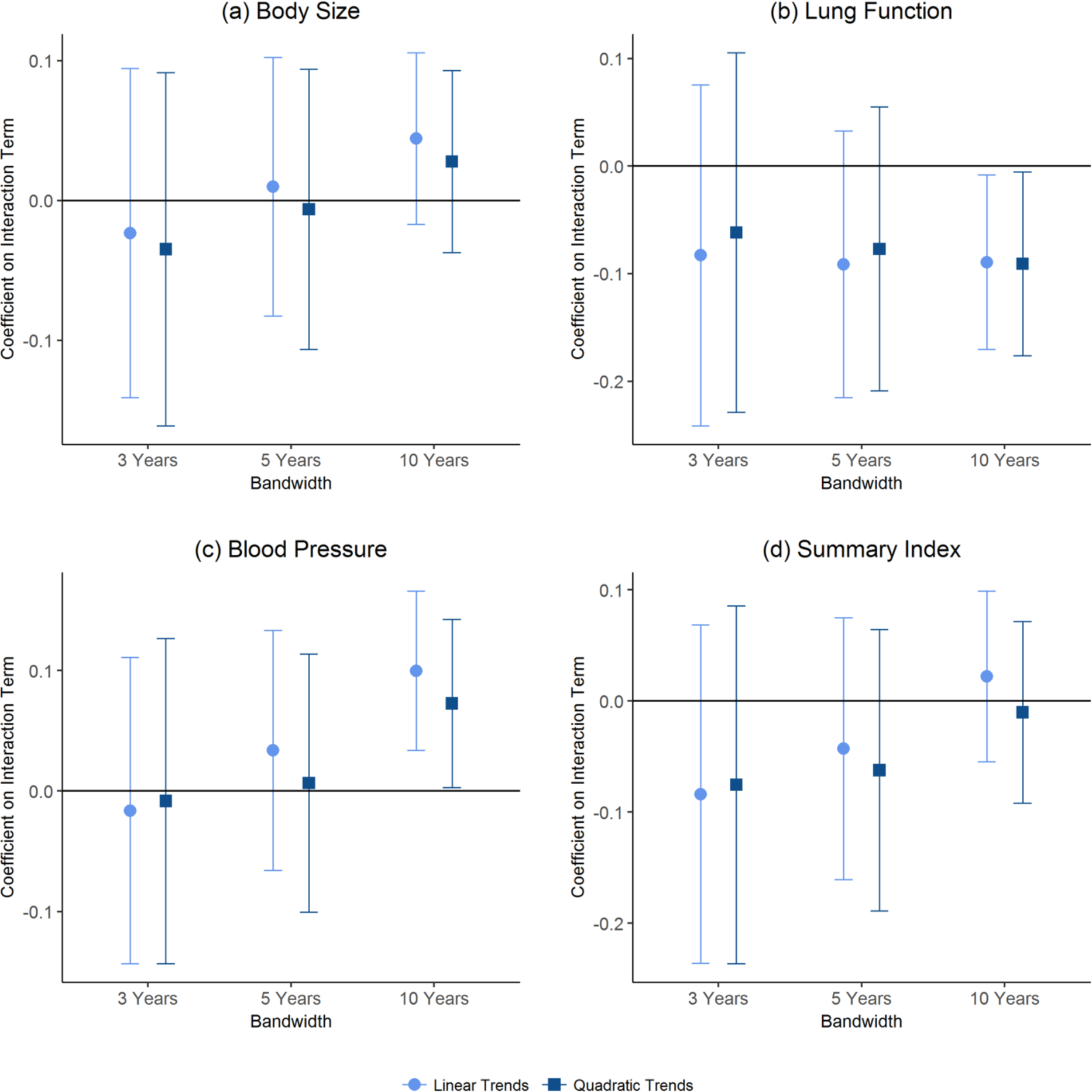
Interaction Estimates for Alternative Bandwidth and Trend Specifications (Continuous Health Indices and BMI PGS). Markers show 2SLS estimates of the coefficient on the interaction of BMI PGS and an indicator for staying in school until age 16 for different bandwidths and birth cohort trends. The dependent variables are continuous measures of (**a**) body size (**b**) lung function (**c**) blood pressure and (**d**) summary indices. The regressions include a quadratic polynomial in date of birth, which is allowed to be different before and after September 1, 1957; the first 15 principal components of the genotypic data; the interactions of those principal components with “Edu 16”; and controls for age, age in days and age squared, dummies for calendar month of birth, and dummies for country of birth. Brackets show 95% confidence intervals.

**Appendix Fig. H2.**
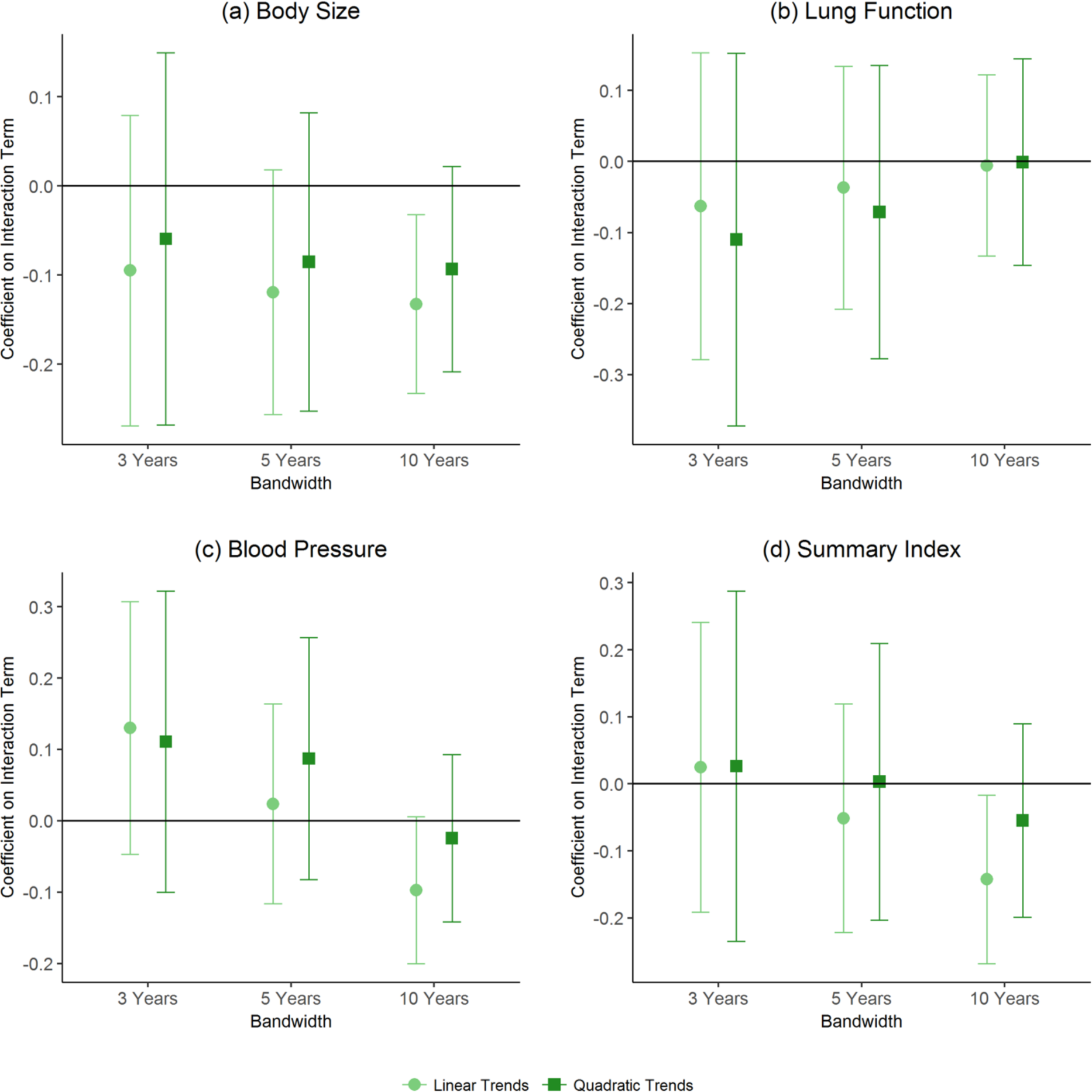
Interaction Estimates for Alternative Bandwidth and Trend Specifications (Continuous Health Indices and EA PGS). Markers show 2SLS estimates of the coefficient on the interaction of EA PGS and an indicator for staying in school until age 16 for different bandwidths and birth cohort trends. The dependent variables are continuous measures of (**a**) body size (**b**) lung function (**c**) blood pressure and (**d**) summary indices. The regressions include a quadratic polynomial in date of birth, which is allowed to be different before and after September 1, 1957; the first 15 principal components of the genotypic data; the interactions of those principal components with “Edu 16”; and controls for age, age in days and age squared, dummies for calendar month of birth, and dummies for country of birth. Brackets show 95% confidence intervals.

**Appendix Fig. H3.**
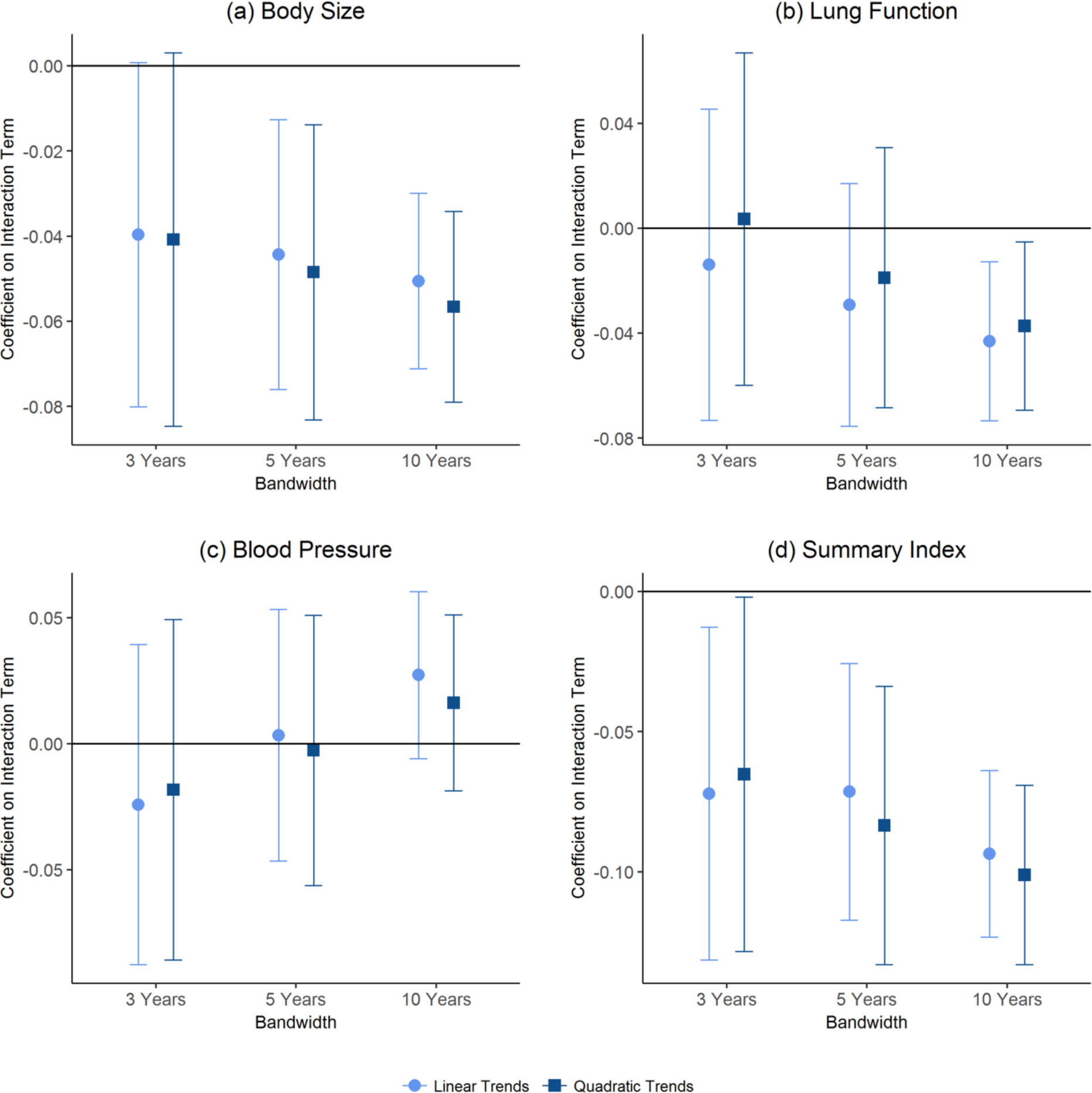
Interaction Estimates for Alternative Bandwidth and Trend Specifications (Binary Health Indices and BMI PGS). Markers show 2SLS estimates of the coefficient on the interaction of BMI PGS and an indicator for staying in school until age 16 for different bandwidths and birth cohort trends. The dependent variables are binary measures of (**a**) body size (**b**) lung function (**c**) blood pressure and (**d**) summary indices. The regressions include a quadratic polynomial in date of birth, which is allowed to be different before and after September 1, 1957; the first 15 principal components of the genotypic data; the interactions of those principal components with “Edu 16”; and controls for age, age in days and age squared, dummies for calendar month of birth, and dummies for country of birth. Brackets show 95% confidence intervals.

**Appendix Fig. H4.**
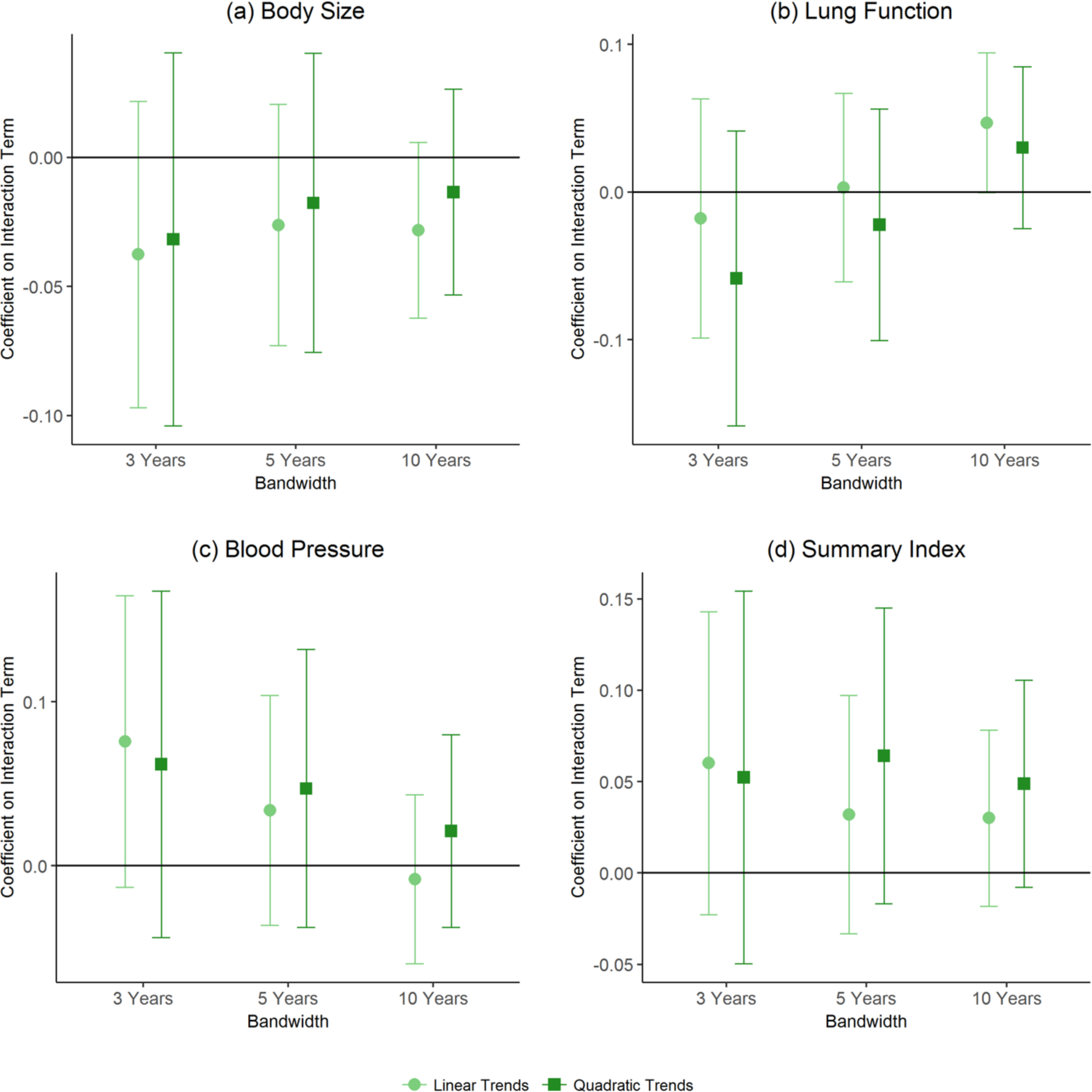
Interaction Estimates for Alternative Bandwidth and Trend Specifications (Binary Health Indices and EA PGS). Markers show 2SLS estimates of the coefficient on the interaction of EA PGS and an indicator for staying in school until age 16 for different bandwidths and birth cohort trends. The dependent variables are binary measures of (**a**) body size (**b**) lung function (**c**) blood pressure and (**d**) summary indices. The regressions include a quadratic polynomial in date of birth, which is allowed to be different before and after September 1, 1957; the first 15 principal components of the genotypic data; the interactions of those principal components with “Edu 16”; and controls for age, age in days and age squared, dummies for calendar month of birth, and dummies for country of birth. Brackets show 95% confidence intervals.

## APPENDIX I: Robustness to Binary Outcomes’ Thresholds

The figures in this section investigate the sensitivity of our results to using different thresholds across the distributions of the indices.

**Appendix Fig. I1.**
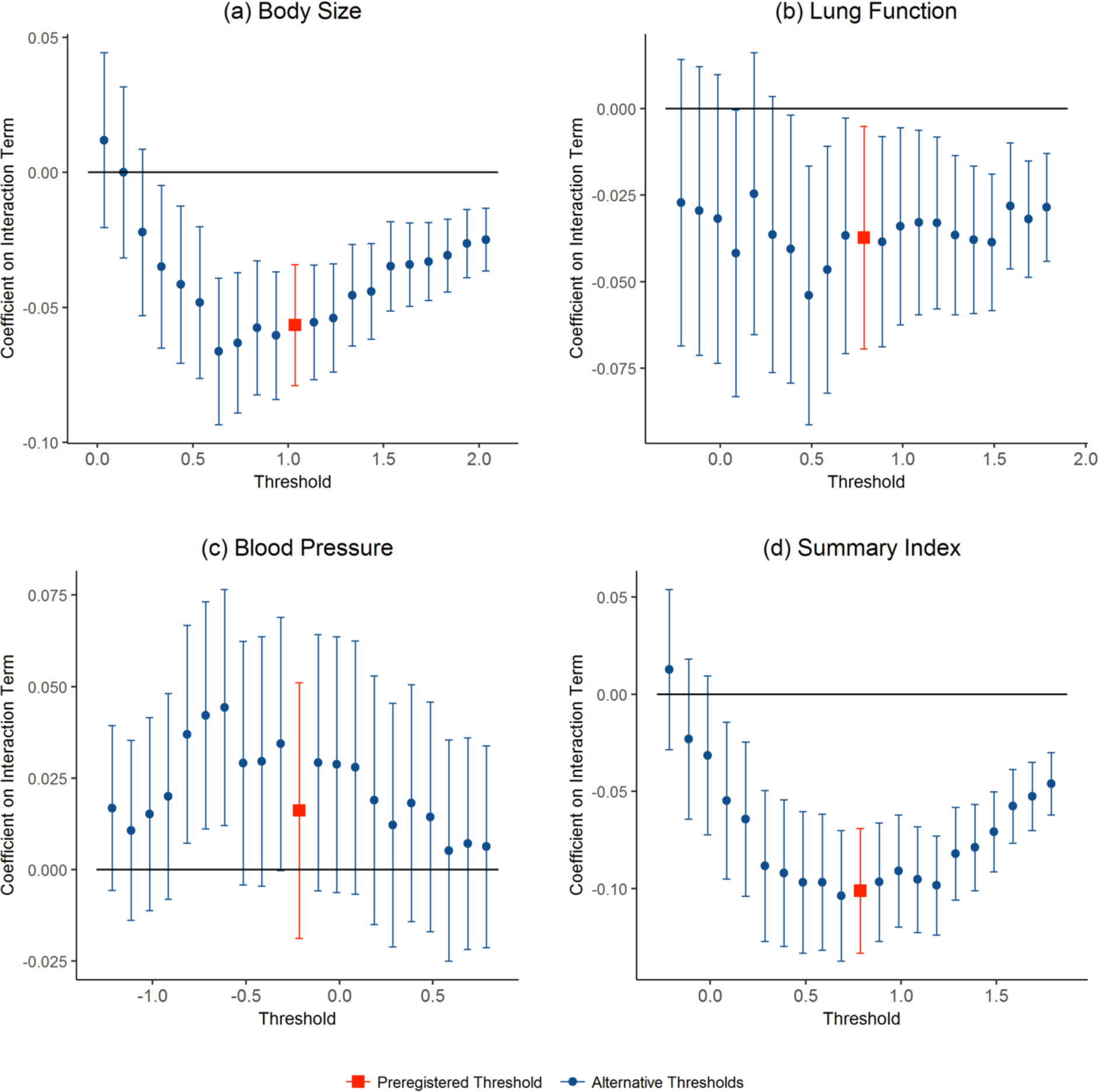
Robustness of Interaction Coefficient Estimates when Alternative Threshold Is Used to Define Binary Health Index (BMI PGS). The figures show the robustness of the coefficient on the BMI PGS and Edu16 interaction to changing the threshold used in the binary measures of (**a**) body size, (**b**) lung function, (**c**) blood pressure, and (**d**) summary indices. Edu 16 is an indicator for staying in school until at least age 16. The red square marker shows estimates of the preregistered threshold. The blue circle markers show estimates for alternative thresholds. The regressions include a quadratic polynomial in date of birth, which is allowed to be different before and after September 1, 1957; the first 15 principal components of the genotypic data; the interactions of those principal components with “Edu 16”; and controls for age, age in days and age squared, dummies for calendar month of birth, and dummies for country of birth. 95% confidence intervals.

**Appendix Fig. I2.**
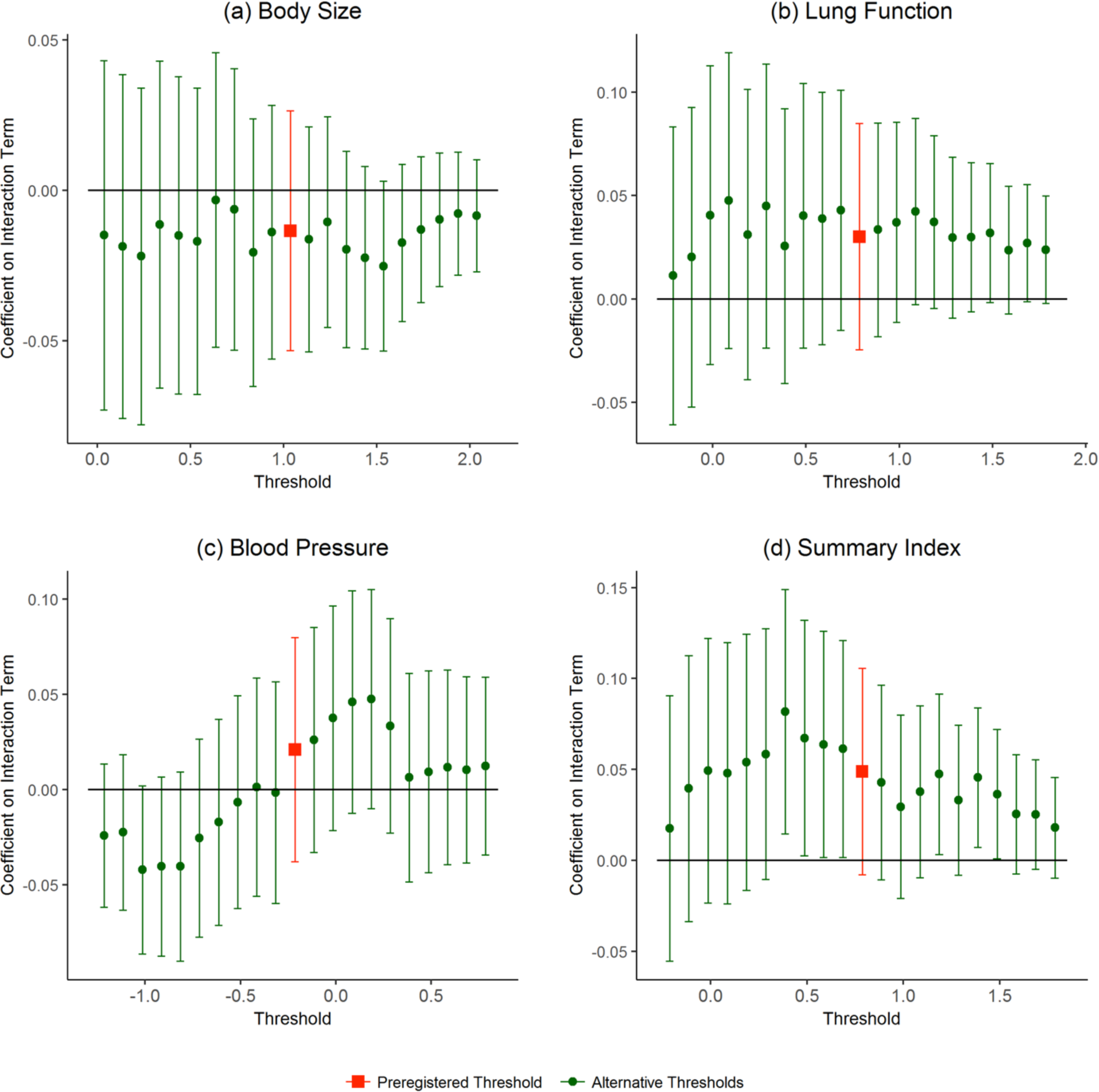
Robustness of Interaction Coefficient Estimates when Alternative Threshold Is Used to Define Binary Health Index (EA PGS). The figures show the robustness of the coefficient on the EA PGS and Edu16 interaction to changing the threshold used in the binary measures of (**a**) body size, (**b**) lung function, (**c**) blood pressure, and (**d**) summary indices. Edu 16 is an indicator for staying in school until at least age 16. The red square marker shows estimates of the preregistered threshold. The blue circle markers show estimates for alternative thresholds. The regressions include a quadratic polynomial in date of birth, which is allowed to be different before and after September 1, 1957; the first 15 principal components of the genotypic data; the interactions of those principal components with “Edu 16”; and controls for age, age in days and age squared, dummies for calendar month of birth, and dummies for country of birth. 95% confidence intervals.

## APPENDIX J: Biomedical Cutoffs

**Appendix Fig. J1.**
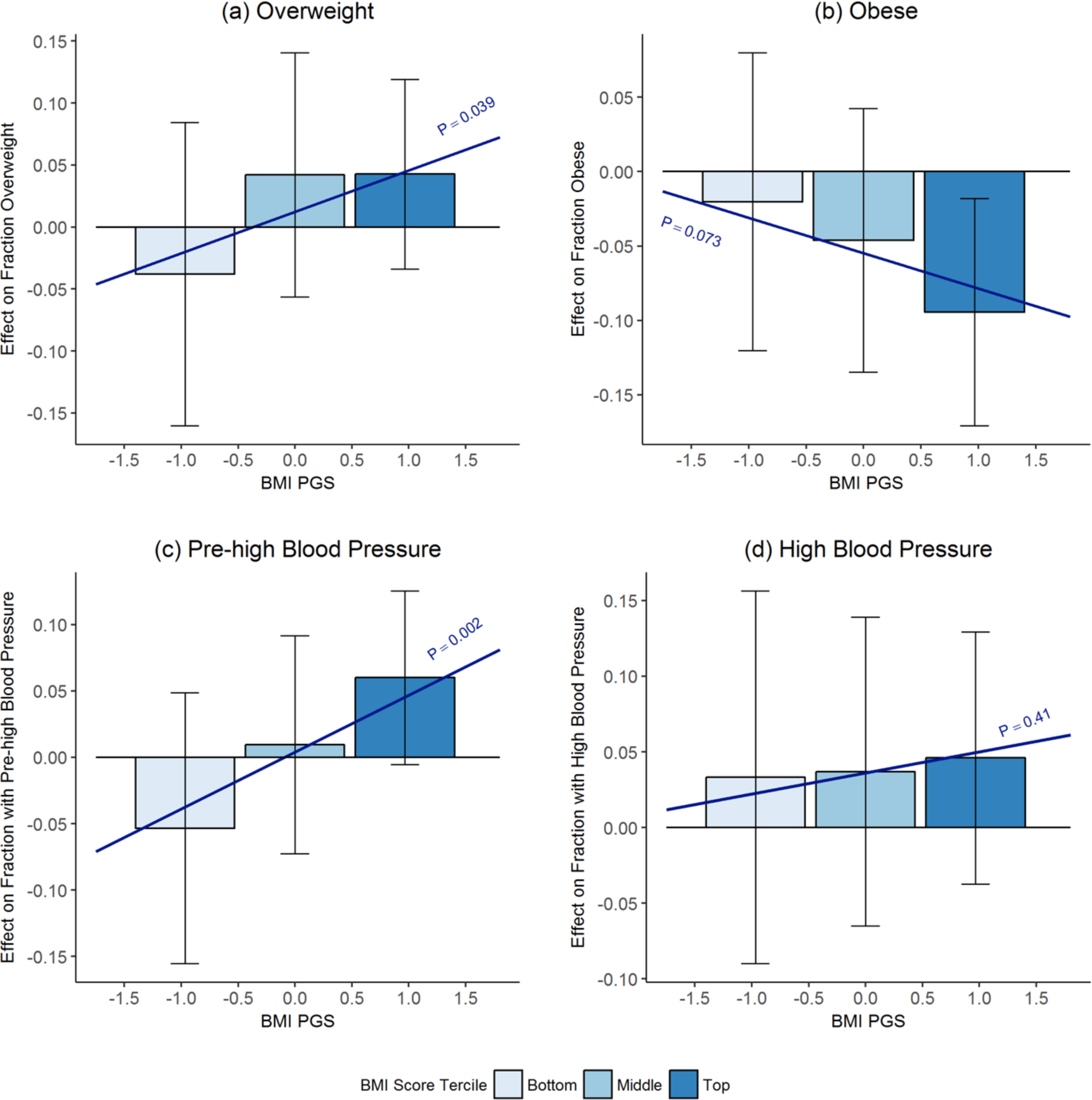
Biomedical cutoffs and BMI PGS. Bars show 2SLS point estimates of effect of staying in school until age 16 on (**a**) fraction overweight (**b**) fraction obese (**c**) fraction with pre-high blood pressure and (**d**) fraction with high blood pressure for the bottom, middle, and top terciles of the BMI PGS distribution. Overweight is defined as having a BMI over 25. Obesity is defined as having a BMI over 30. Pre-high blood pressure is defined as having diastolic blood pressure above 80 or systolic blood pressure above 120. High blood pressure is defined as having diastolic blood pressure above 90 or systolic blood pressure above 140. Brackets show 95% confidence intervals. Sloped lines plot 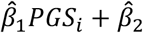. “P” corresponds to the p-value of *H*_0_: β_1_ = 0.

**Appendix Fig. J1.**
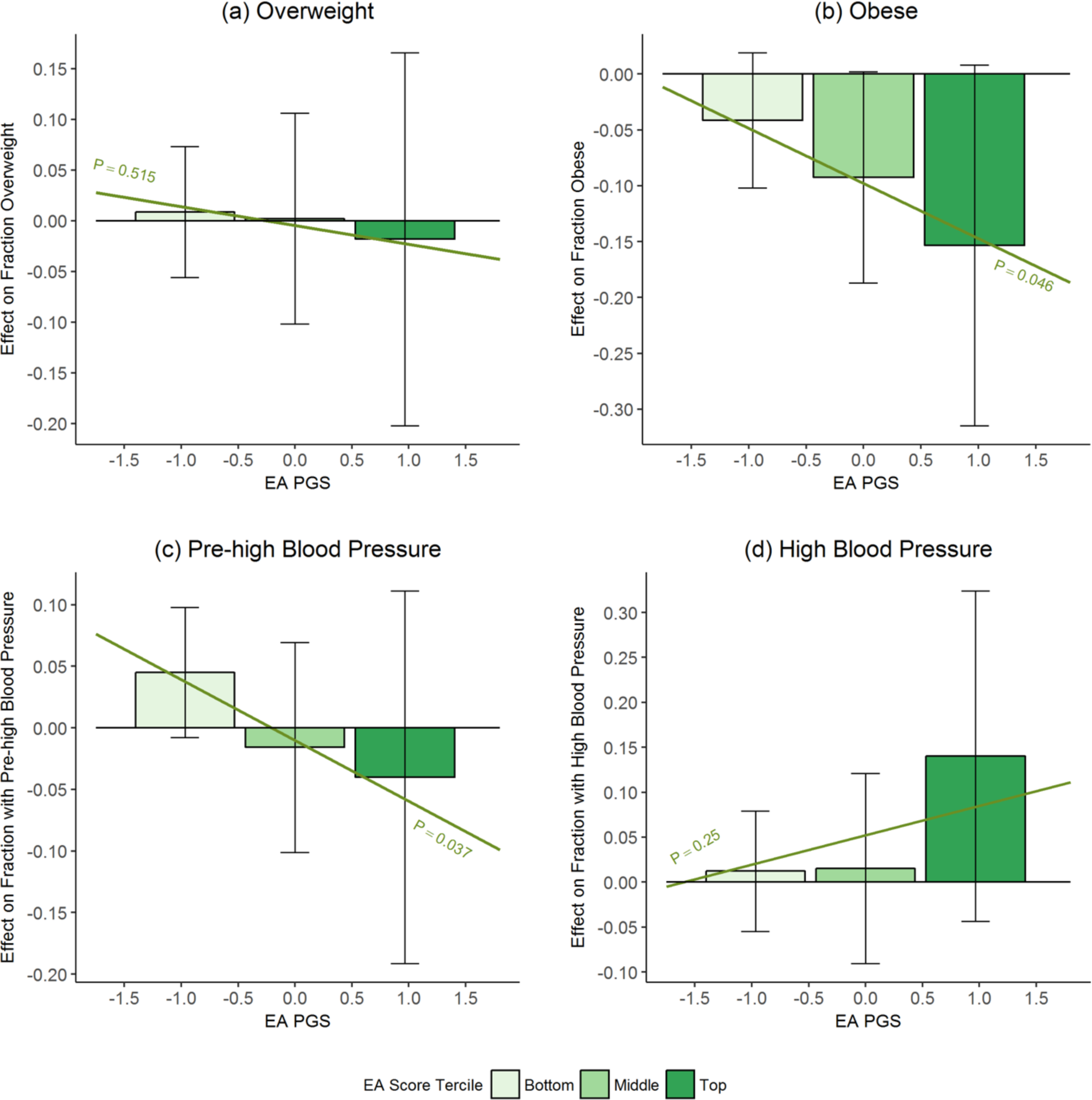
Biomedical cutoffs and EA PGS. Bars show 2SLS point estimates of effect of staying in school until age 16 on (a) fraction overweight (b) fraction obese (c) fraction with pre-high blood pressure and (d) fraction with high blood pressure for the bottom, middle, and top terciles of the EA PGS distribution. Overweight is defined as having a BMI over 25. Obesity is defined as having a BMI over 30. Pre-high blood pressure is defined as having diastolic blood pressure above 80 or systolic blood pressure above 120. High blood pressure is defined as having diastolic blood pressure above 90 or systolic blood pressure above 140. Brackets show 95% confidence intervals. Sloped lines plot 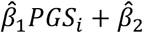. “P” corresponds to the p- value of *H_0_: β_1_* = 0.

## APPENDIX K: Mechanisms

**Appendix Table K1.**
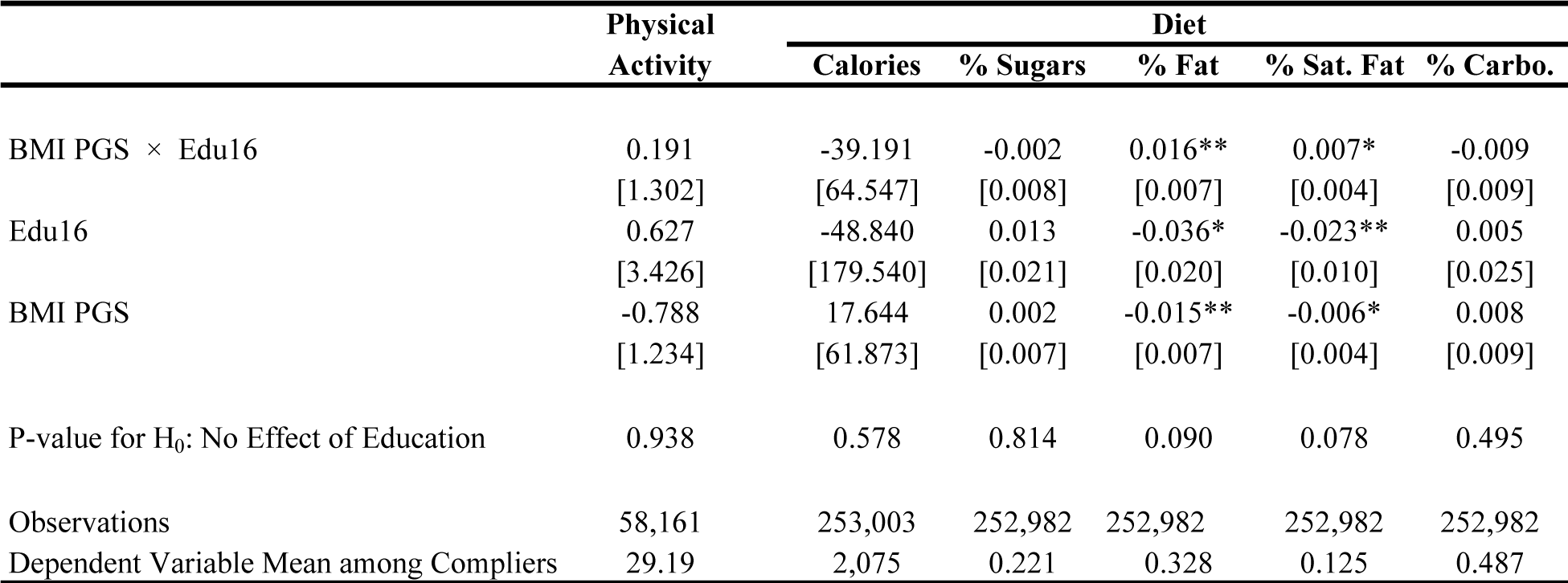
Effect of staying in school until age 16 on obeseity-related health behaviors. 2SLS estimates. Edu16 is an indicator for staying in school until age 16 and is instrumented by an indicator for being born after September 1, 1957. The “P-value for H_0_: No Effect of Education” is the p-value from a joint test that the coefficient on “BMI PGS x Edu16” and on “Edu16” are each equal to zero. The last row shows means of the dependent variable among pre-reform compliers, defined as individuals born before September 1, 1957 who dropped out before age 16.

## Footnotes

1 The NHS has contact details for an estimated 98% of the UK population.

2 The UK Biobank provides two measures of BMI: one calculated as weight in kilograms divided by height squared (in meters) and one using height and electrical impedance to quantify mass. We will take the average of these two measures. The waist-hip ratio is equal to the waist circumference divided by the hip circumference.

3 They were instructed to continue blowing until no more air came out of their lungs. Up to three attempts were allowed. The participant was allowed a third attempt if the first two blows did not satisfy the reproducibility criteria of the spirometry protocol.

4 Principal components are included to control for ancestry effects that may be correlated with a person’s polygenic score (Price et al. 2006). Doing so is standard in analyses using genome-wide data.

5 Gelman and Imbens (2016) caution against the use of higher order polynomials (higher than 2) in RD.

6 Because participants were surveyed for the baseline assessment between 2006 and 2010, date of birth and age are not perfectly collinear.

